# E-cadherin/HMR-1 and PAR-3 break symmetry at stable cell contacts in a developing epithelium

**DOI:** 10.1101/2022.08.10.503536

**Authors:** Victor F. Naturale, Melissa A. Pickett, Jessica L. Feldman

## Abstract

Tissue-wide patterning is essential to multicellular development, requiring cells to individually generate polarity axes and coordinate them in space and time with neighbors. Using the *C. elegans* intestinal epithelium, we identified a patterning mechanism informed by stabilized cell/cell contact and executed via the scaffolding protein PAR-3 and the transmembrane protein E-cadherin/HMR-1. Intestinal cells break symmetry as PAR-3 and HMR-1 recruit apical determinants into micron-scale ‘local polarity complexes’ (LPCs) at homotypic contacts. LPCs undergo a HMR-1-based migration to a common tissue midline, thereby establishing tissue-wide polarity. Thus, symmetry breaking results from PAR-3-dependent intracellular polarization coupled to HMR-1-based tissue-level communication that occurs through a non-adhesive signaling role for HMR-1. Intestinal cells gain initial asymmetry from differential contact duration as homotypic contacts last longer than heterotypic contacts, thus providing stable platforms for LPC assembly and offering a logical and likely conserved framework for how internal epithelia with no obvious pre-existing asymmetries can polarize.

## Introduction

Cell polarity is the asymmetric separation of molecules into distinct subcellular domains, which underlies many biological processes and allows cells to execute diverse functions. Fundamentally, polarizing systems only require molecules that positively reinforce one another and differ in diffusion rate (Gierer and Meinhardt, 1972; Turing, 1952; Vendel et al., 2019). To impart biological functions, these properties are often linked to extrinsic cues, allowing cells to ‘break symmetry’ and define polarity axes in response to their environment. In some situations, these cues are required to initiate polarity, while in others they simply orient an inherently self-segregating set of molecules. For example, in the one-cell *C. elegans* embryo, a sperm entry cue is needed to establish antagonizing PAR domains to define the anterior/posterior body axis (Munro and Bowerman, 2009). In contrast, lipids within *Dictyostelium* cell membranes stochastically self-organize in the absence of external cues but are sharply oriented in response to chemoattractants (Shibata et al., 2013).

Within multicellular organisms, cells often polarize collectively to build tissue-wide patterns. These contexts present an added challenge, as symmetry breaking must occur not only within every cell but also coordinately between neighbors to generate a common polarity axis. Thus, global cues likely exist to ensure reproducible symmetry breaking across whole tissues. This problem is clearly exemplified in epithelia, adherent sheets of cells which encase the internal organs of metazoans. Epithelia protect underlying tissues from the external environment while also possessing context-specific functions that depend on establishing polarized apical and basolateral membrane domains. Most apical domains localize the conserved PAR polarity complex – scaffolds Par3/PAR-3 and Par6/PAR-6, and the kinase aPKC/PKC-3 – and orient toward contact-free surfaces or into a hollow lumen. Basolateral domains localize separate components like the Scribble complex to surfaces contacting neighboring epithelial cells or underlying tissues (Pickett et al., 2019). Separating apical and basolateral domains are junctional complexes including adherens junctions, which contain the transmembrane adhesion molecule E-cadherin and associated adaptors (Vasquez et al., 2021). In minimal systems, several of these polarity proteins are known to self-organize into discrete domains owing to their capacity for positive feedback and mutual antagonism (Kono et al., 2019). However, epithelial polarization must occur with precise timing and reproducible orientation during development, as delays or errors in this process can lead to embryonic lethality and/or congenital defects (Cui et al., 2022; Mescher et al., 2017; Mizotani et al., 2018; Schneeberger et al., 2018; Tanentzapf et al., 2000; Totong et al., 2007). Thus, epithelial symmetry breaking at cell and tissue scales cannot rely on self-organization alone and must involve other orienting cues.

Although epithelia develop in many unique contexts and have evolved equally varied polarity programs (Pickett et al., 2019), some common symmetry breaking themes have emerged. For example, many epithelia orient their polarity in reference to an underlying basement membrane, as integrin receptors bind to ECM and facilitate intracellular apical exclusion from these contacts (Myllymäki et al., 2011; O’Brien et al., 2001; Rasmussen et al., 2012; Yu et al., 2005). Mitosis can also orient polarity, as the midbody remnant left behind from cell division can act as an Apical Membrane Initiation Site (‘AMIS’) to direct trafficking of apical cargo (Bryant et al., 2010, 2014; Li et al., 2014; Lujan et al., 2017). Finally, cell/cell contact between epithelial neighbors can provide polarity information. When devoid of any contact, individual epithelial cells fail to polarize (Rodriguez-Boulan et al., 1983; Wang et al., 1990; Yeaman et al., 1999). They can be genetically induced to do so upon overactivation of polarity machinery (Baas et al., 2004), but resulting apical domains within cells fail to orient in a common direction in the absence of contact. Many studies in epithelial and non-epithelial cells have explored E-cadherin as a link between cell/cell contact and apical domain orientation. Indeed, in some contexts E-cadherin is sufficient to instruct aspects of polarity (Cohen et al., 2007; Klompstra et al., 2015; Liang et al., 2022; Zhang et al., 2020). However, specific symmetry breaking functions for E-cadherin have often been obscured by its essential role in cell/cell adhesion (Cox et al., 1996; Johnson et al., 1986; Peifer et al., 1993; Shirayoshi et al., 1983; Stephenson et al., 2010). Moreover, many insights into epithelial symmetry breaking come from cultured cell lines which lack the developmental contexts in which polarity normally occurs, leaving open the question of how the presence of and response to symmetry breaking cues may be temporally regulated in coordination with the rest of an organism.

Here, we use the *C. elegans* embryonic intestine to understand cell- and tissue-level symmetry breaking *in vivo.* The *C. elegans* intestine is a simple epithelial tube comprised of 20 ‘E’ cells. Intestinal cells are clonally derived from a single blastomere (E) and adopt apico-basolateral polarity by the 16-cell (‘E16’) stage, with all apical surfaces oriented toward a common embryonic midline that will give rise to the future lumen and with basolateral domains at remaining membranes (Fig 1A, 1E; Leung et al., 1999). Importantly, intestinal polarization occurs independent of a known basement membrane (Graham et al., 1997; Rasmussen et al., 2012) and at sites distinct from those where midbodies are positioned, so neither of these well-known cues likely instructs symmetry breaking in this context. Indeed, the E blastomere can develop *ex vivo* into a polarized cyst (Leung et al., 1999), showing that symmetry breaking information is contained within the primordium and independent of tissue-extrinsic cues.

**Figure 1.**
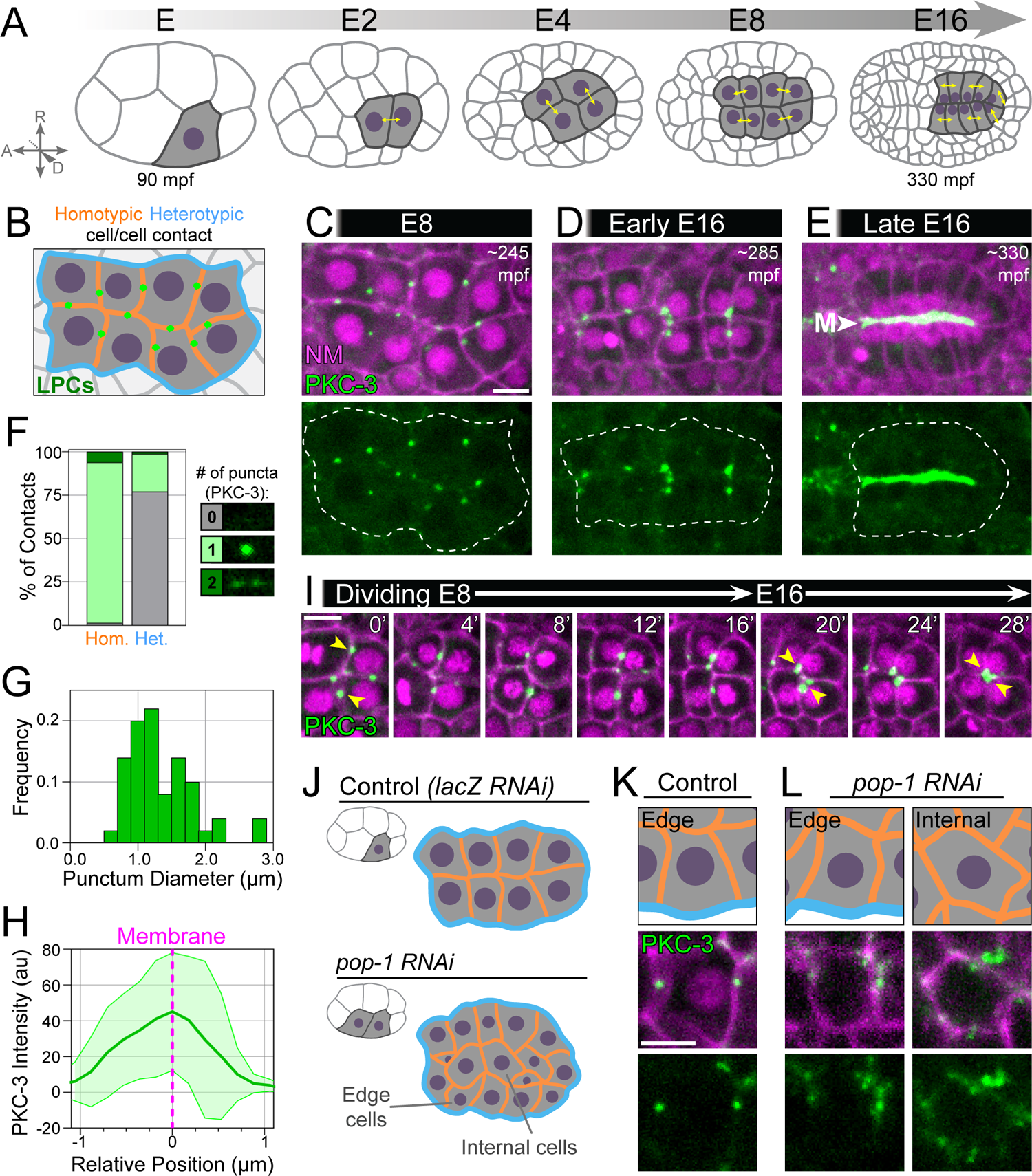
The *C. elegans* intestine breaks symmetry at homotypic contacts through discrete cell- and tissue-level steps. A) Dorsal view cartoon of intestinal development from E to E16 stage with sister cells indicated (yellow double arrow; mpf = minutes post fertilization). B) Cartoon of homotypic/heterotypic contact pattern at E8, LPCs in green. C-E) Cropped dorsal views of live E8, ‘Early’ E16 (0’ post division), and ‘Late’ E16 (∼40’ post division) intestines (white dotted lines). NM = histone::mCherry, membrane::mCherry, M = intestinal midline. F) Number of PKC-3 puncta at homotypic vs heterotypic contacts in E8-stage (∼245-265 mpf) embryos (n=65 homotypic and 78 heterotypic contacts, 13 embryos. G) Distribution of E8 PKC-3 puncta size (n=64 puncta, 13 embryos). H) Intensity of PKC-3 relative to membrane (magenta dotted line). Lines and shading represent mean +/- SD (n=64 puncta, 13 embryos). I) Cropped (dorsal view) timelapse of 4 E8 cells. Yellow arrowheads highlight LPC retention and migration. J) Cartoon of E contacts in embryos treated with lacZ or *pop-1* RNAi, with 8-cell embryo cartoon (top left) depicting MS to E fate conversion. K-L) Live imaging of ‘edge’ or ‘internal’ E cells from embryos treated with lacZ or *pop-1* RNAi (lacZ: n=10 embryos; *pop-1*: n=8 embryos). Scale bars=5 μm.

We report a two-step intrinsic symmetry breaking mechanism reliant on homotypic (E cell/E cell) contact. E cells initially break symmetry through the assembly of ‘local polarity complexes’ (LPCs), large punctate structures of apical and junctional proteins, at homotypic contacts between intestinal precursors. The intestinal primordium then breaks symmetry at the tissue level as LPCs are shuttled along membranes to the intestinal midline to generate a continuous apical surface. These choreographed movements are controlled by the polarity scaffold PAR-3 and the transmembrane protein E-cadherin/HMR-1, which act in parallel to build LPCs but perform separable functions to enlarge and then move LPCs during tissue-level symmetry breaking. Importantly, this symmetry breaking function of HMR-1 occurs separately from its role in general adhesion and at the adherens junction. Finally, we demonstrate that the information used to pattern LPCs is derived from differences in contact stability, as homotypic contacts are inherently longer lived than heterotypic contacts and provide more time for LPC formation. Together, these data identify differential contact lifetimes as a symmetry breaking cue within a developing epithelium and demonstrates how this cue is translated into cell- and tissue-level polarity through a PAR-3 and E-cadherin-based symmetry breaking module.

## Results

### The *C. elegans* intestine breaks symmetry at homotypic contacts through discrete cell- and tissue-level steps

We first sought to identify when and how endogenous PAR proteins localize asymmetrically within intestinal cells prior to reaching the midline, the future apical surface of this tissue. Consistent with previous reports, PAR complex members PAR-3, PAR-6, and PKC-3 localized within discrete puncta at E8-stage cell/cell contacts, one cell cycle before PAR-containing puncta coalesce at the midline at E16 (Fig. 1B-D; Fig. S1; Achilleos et al., 2010; Feldman and Priess, 2012). To reflect the localization and composition of these structures, we hereafter refer to them as Local Polarity Complexes (LPCs). LPCs localized almost exclusively to E/E ‘homotypic’ contacts and were excluded from E/non-E ‘heterotypic’ contacts (Fig. 1B, 1F). 92% of homotypic contacts possessed a single LPC (mean diameter=1.35 μm; Fig. 1F, 1G), which generally localized midway along the length of each contact (Fig. S1C). LPCs appeared to localize on the membrane and associate with both cell neighbors, as (1) PKC-3 intensity at LPCs peaked at the membrane (Fig. 1H) and (2) nuclei from multiple cells at later stages apposed LPCs and clustered around them (Fig. 1D; Feldman and Priess, 2012). Thus, LPCs are stereotyped and actively positioned structures which exist over one hour before tissue-level polarization.

The apical surfaces of intestinal cells are established at E16 as puncta of PAR proteins localized on lateral membranes of adjacent non-sister cells migrate toward and spread along the midline over 30-40 minutes (Fig. S1A; Achilleos et al., 2010; Feldman and Priess, 2012). We therefore wanted to understand whether the LPCs present at E8 are the same puncta that have been observed at E16. LPCs remained on homotypic contacts throughout E8, even as some cells repositioned themselves within the primordium to achieve a bilayered arrangement prior to division (Fig. S1B; Asan et al., 2016). As E8 cells divided, some LPCs were retained at contacts (Fig. 1D, 1I). When mitosis completed, LPCs underwent a directed movement to the midline before spreading across it to establish the apical surface (Fig. 1E, 1I; Movie S1). Thus, apical PAR proteins first localize to stereotyped LPCs, which act as seeds as they are retained through morphogenesis and migrate to the midline to establish a common apical surface. We therefore define symmetry breaking in this system as occurring in two separable steps: 1) cellular symmetry breaking, when local E neighbors package PAR proteins in LPCs at E8; and 2) tissue symmetry breaking, when LPCs migrate to a common surface within the E16 primordium, the intestinal midline.

### Homotypic contact informs PAR protein localization

Since LPCs first localize to homotypic contacts during cellular symmetry breaking, we next wanted to determine whether E/E contact plays an instructive role in PAR protein localization. To test this hypothesis, we asked whether LPCs form at ectopic E contact sites. We treated embryos with RNAi against the Wnt pathway component *pop-1*/TCF, a fate determinant required to suppress E fate in the MS blastomere (Fig. 1J; Lin et al., 1995). *pop-1* RNAi transformed the MS lineage into a second E lineage; these embryos contained about twice the number of E cells, consistent with previous reports, and these cells were often packed into extra rows in the primordium (Fig. 1J). As in control embryos, cells at the borders of the intestinal primordium (‘edge cells’; Fig. 1J) within *pop-1* RNAi-treated embryos localized PKC-3 specifically to homotypic contacts and not to heterotypic contacts (Fig. 1K, 1L left panel). In contrast, where E cells were surrounded by other E neighbors (‘internal cells’; Fig. 1J), PKC-3 localized to all contacts (Fig. 1L, right panel). These results demonstrate that altered PAR localization is a result of changes in contact rather than *pop-1* depletion per se, implicating a mechanism of cellular symmetry breaking informed by homotypic cell/cell contacts.

### LPCs are born from small foci and mature in composition and structure over time

We next wanted to understand the composition and genesis of the LPC, initially focusing on PAR-3 as the most upstream known polarity protein in this system (Achilleos et al., 2010). The E4 stage was the earliest timepoint that we observed endogenous PAR-3 at intestinal cell/cell contacts (Fig. 2A-B), when it localized to small, dim ‘foci’ rather than the large punctate LPCs seen at E8 (Fig. 2B, inset vs Fig. 2B’’’, inset). E4 foci were ephemeral, and contacts often contained multiple foci which appeared and disappeared over time in no clear pattern. Foci were enriched at homotypic contacts but also existed at heterotypic contacts and at three- and four-cell vertices (Fig. 2B and data not shown). Upon entry into E8, PAR-3 foci became fewer but longer-lasting and brightened into larger LPCs (Fig. 2B’-2B’’’). Heterotypic PAR-3 foci were still identifiable at E8, although they were dimmer and more ephemeral than homotypic LPCs. Surprisingly, PAR-3 foci did not initially co-localize with other apical PAR proteins PAR-6 and PKC-3, which were instead diffuse along the cell cortex and in the cytoplasm at E4 and did not enrich in LPCs until after the start of E8 (Fig. 2A, ‘PAR Complex’; Fig. 2C-2C’’’, Fig. S1D). In addition, PKC-3 accumulated more gradually into one LPC per contact rather than localizing first to multiple foci, suggesting that PAR-3 and PKC-3 assemble into LPCs in distinct ways.

**Figure 2.**
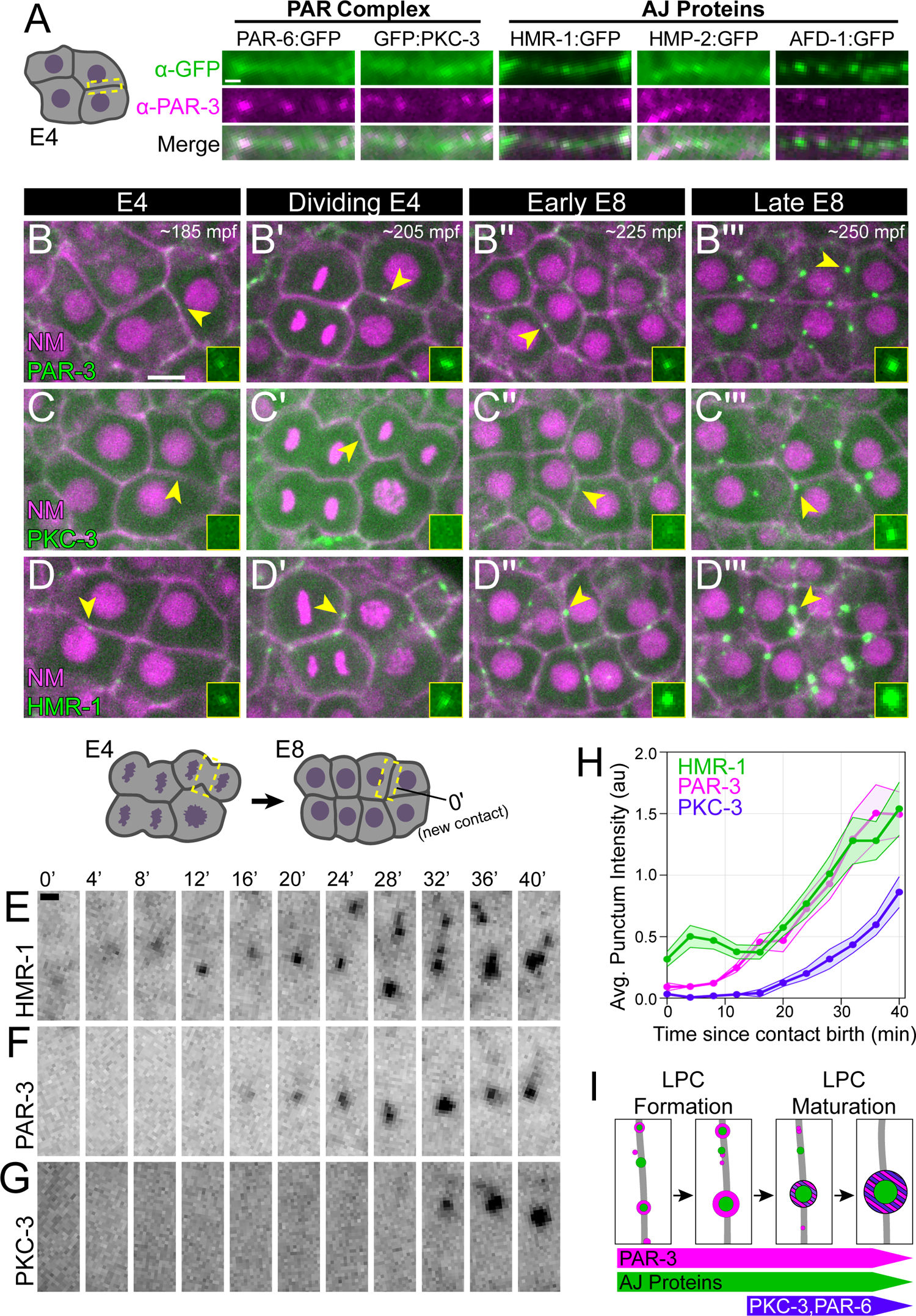
LPCs are born from small foci and mature in composition and structure over time. A) Left: Cartoon of E4 intestine with indicated homotypic contact shown at right (yellow dotted box). Right: α-GFP, α-PAR-3 immunostaining of fixed E4 embryos represented as single slices and expressing indicated endogenous GFP-tagged proteins. B-D’’’) Live imaging of indicated endogenously GFP-tagged protein in dorsal view E4-E8 stage intestine (NM = histone::mCherry, membrane::mCherry). Yellow arrowheads point to GFP signal highlighted in insets. E-G) Top: Cartoon of E4-E8 division with region shown at bottom indicated (dotted yellow box). Bottom: Stage-matched time courses of indicated endogenously GFP-tagged protein. H) Analysis of HMR-1, PAR-3, and PKC-3 intensity over time. Lines and shading represent mean +/- SEM (PAR-3: n=24 contacts, 6 embryos; HMR-1: n=24 contacts, 6 embryos; PKC-3: n=12 contacts, 3 embryos). I) Cartoon summary of LPC formation and maturation. Scale bar=1 μm (A, E-G) or 5 μm (B-D’’’, insets magnified 2x).

PAR proteins co-localize and co-migrate with adherens junction (AJ) proteins at E16 (Pickett et al., 2021; Totong et al., 2007). Consistently, E-cadherin/HMR-1, β-catenin/HMP-2, α-catenin/HMP-1, and afadin/AFD-1 were all found in E4 foci and sometimes colocalized with or were adjacent to PAR-3 before enriching together into LPCs at the E8 stage (Fig. 2A, ‘AJ Proteins’; Fig. S1D). AJ proteins behaved similarly to PAR-3 through development, as HMR-1 also localized to short-lived foci before enriching into LPCs with PAR-3 (Fig. 2D-2D’’’).

To precisely map a timeline of LPC genesis, we examined PAR-3, HMR-1, and PKC-3 at a newly born E8-stage contact (Fig. 2E-H). HMR-1 was first to appear at membranes although initially was quite diffuse (Fig. 2E, t=0’-8’). Within 12-16 minutes, HMR-1 condensed into foci concomitant with the appearance of dim PAR-3 foci (Fig. 2F). PAR-3 and HMR-1 intensity then grew linearly and clusters of smaller foci merged or dissipated yielding a single brighter LPC (Fig. 2E, 24’-40’; Fig 2H). In contrast to PAR-3 and HMR-1, PKC-3 fluorescence was dim until 28-32 minutes after contact birth (Fig. 2G-H, ∼16 minutes after HMR-1/PAR-3 localization) before also increasing in intensity (Fig. 2H). Thus, LPCs are born from the consolidation of HMR-1 and PAR-3 foci at cell/cell contacts into more stable nascent LPCs (‘Formation’, Fig. 2I) which mature when they localize PKC-3 and presumably its obligate binding partner PAR-6 (‘Maturation’, Fig. 2I; Fig. S1D).

### An interdependent HMR-1/PAR-3 module is necessary for cellular symmetry breaking

This sequence of LPC genesis suggested that HMR-1 or PAR-3 may be required for LPCs formation. We tested this hypothesis using an intestine-specific protein depletion strategy: HMR-1 and PAR-3 were endogenously tagged with a ZF::GFP degron, and degradation was achieved through expression of the E3 ligase component ZIF-1, which directs degradation of ZF-tagged proteins, under the intestinal *elt-2* promoter driving robust degradation by E8 (Armenti et al., 2014; Sallee et al., 2018; Fig. S2A, S2D). LPCs were examined in each depletion at the end of E8 (∼260mpf), when PKC-3 became robustly localized to LPCs in control embryos (Fig. 3A). Embryos with intestine-specific PAR-3 depletion (‘PAR-3^gut(-)^’) lacked PKC-3 puncta (Fig. 3A’, 3B), consistent with the known role of PAR-3 to recruit PKC-3 and PAR-6 (Achilleos et al., 2010). HMR-1^gut(-)^ embryos also lacked most or all PKC-3 localization (Fig. 3A’’, 3B), with some smaller and dimmer PKC-3 puncta occasionally visible (remaining puncta were 58% smaller, p=0.0010, and 64% dimmer, p=0.0006, than control). In contrast, HMR-1 localization to LPCs was still strong in PKC-3^gut(-)^ embryos (Fig. 3D vs Fig. 3D’’). Thus, HMR-1 and PAR-3 are both required upstream of PKC-3 localization and therefore LPC maturation.

**Figure 3.**
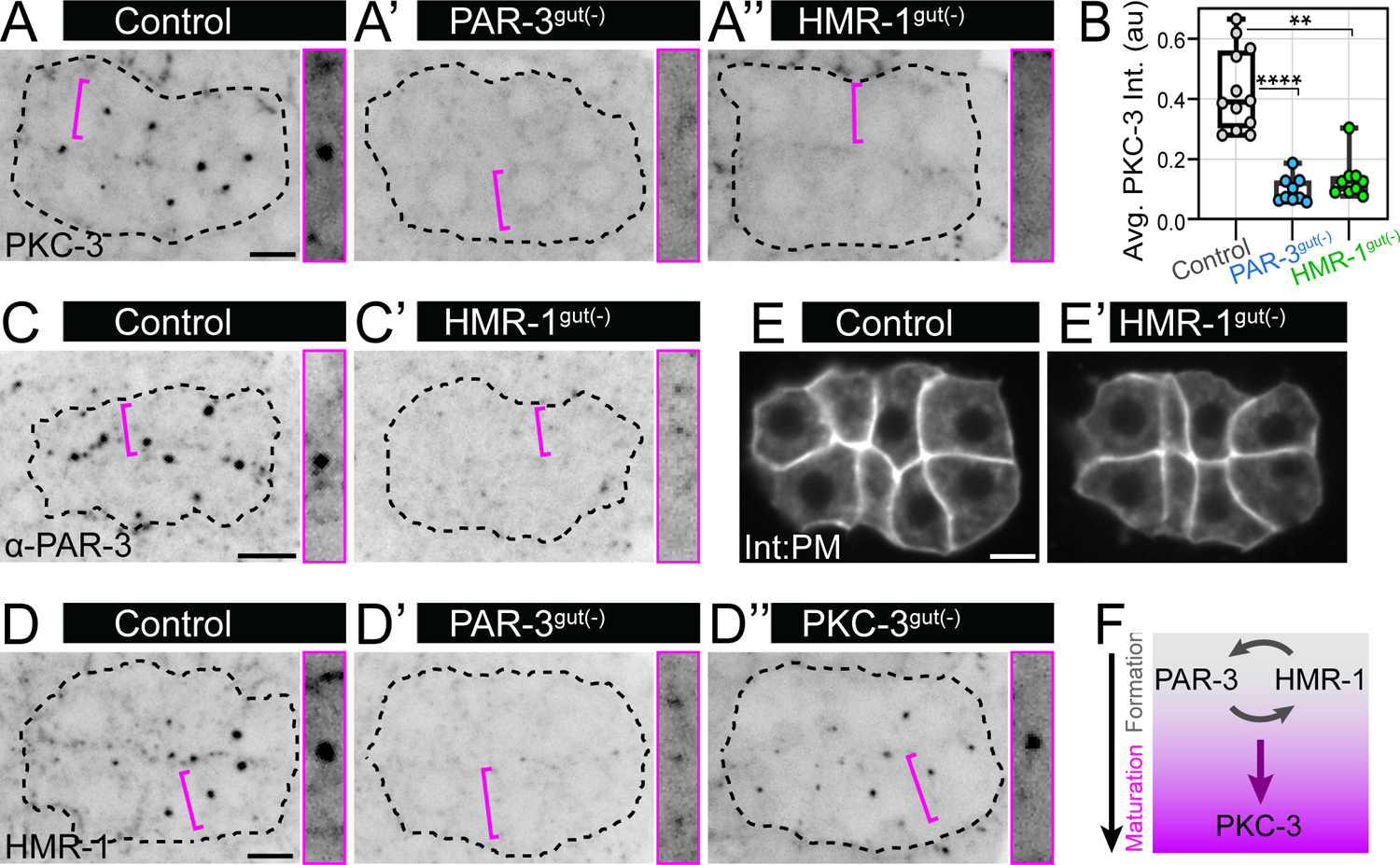
An interdependent HMR-1/PAR-3 module is necessary for cellular symmetry breaking. Dorsal views of E8 intestinal primordia (∼260 mpf) in live embryos showing the indicated endogenously tagged protein (A-A’’’, D-D’’’) or fixed embryos stained with α-PAR-3 antibody (C-C’). Magenta insets at right show one contact at high contrast. Intestines outlined in black. A-A’’) GFP:PKC-3 localization in live Control (n=12), PAR-3^gut(-)^ (n=9), or HMR-1^gut(-)^ (n=9) embryos. B) PKC-3 intensity analysis corresponding to images in A-A’’. Each dot represents one embryo. ****p<0.0001, **p<0.01. C-C’) α-PAR-3 staining in fixed Control (n=11) and HMR-1^gut(-)^ (n=13) embryos. D-D’’) HMR-1:GFP in live Control (n=13), PAR-3^gut(-)^ (n=13), or PKC-3^gut(-)^ (n=11) embryos. E-E’) E cell membranes as visualized by *elt-2p::*GFP::CAAX in live Control (n=10) and HMR-1^gut(-)^ (n=17) embryos. F) Cartoon of genetic interactions during LPC assembly. Scale bars=5 μm (insets magnified 1.5x).

We next tested whether HMR-1 and PAR-3 recruit one another to LPCs. In HMR-1^gut(-)^ animals, PAR-3 puncta were dim and diffuse compared to control, but bore a resemblance to the PAR-3 foci normally visible in wildtype E4 cells (Fig. 3C vs 3C’, insets). HMR-1 localized to similar small foci along homotypic contacts upon PAR-3 depletion (Fig. 3D vs 3D’, insets), leading us to speculate that PAR-3 and HMR-1 are competent to self-organize into foci but require positive feedback to enlarge into an LPC. Importantly, loss of PKC-3 and PAR-3 puncta upon HMR-1 depletion was not merely a byproduct of lost cell/cell adhesion, as E cells in HMR-1^gut(-)^ embryos remained in close contact (Fig. 3E,E’) consistent with HMR-1 being dispensable for adhesion in the *C. elegans* embryo (Grana et al., 2010). Together, the onset of cellular symmetry breaking relies on an interdependent module of HMR-1 and PAR-3, which populate and enlarge the LPC and promote the timely recruitment of other apical determinants (Fig. 3F).

### HMR-1 and PAR-3 accomplish tissue-level symmetry breaking through separable roles

We next investigated whether HMR-1 and PAR-3 are required for tissue symmetry breaking, when LPCs migrate and spread along the intestinal midline at E16. We observed HMR-1 localization in PAR-3^gut(-)^ embryos and PKC-3 in HMR-1^gut(-)^ embryos because PAR-3 fluorescence was too dim to reliably track and PKC-3 and PAR-3 always colocalize at this stage (Pickett et al., 2021). By E16, PKC-3 and HMR-1 became brighter in HMR-1^gut(-)^ and PAR-3^gut(-)^ embryos, respectively, although still localizing as small foci rather than large LPCs. PKC-3-marked LPCs migrated to the midline in control embryos, but this migration failed in HMR-1^gut(-)^ embryos as PKC-3 foci instead stalled at lateral membranes and sometimes drifted basally (Fig. 4A-C). In PAR-3^gut(-)^ embryos, some dim HMR-1 foci also stalled relative to control (Fig. 4D, yellow arrowhead), though HMR-1 gradually appeared at the midline and intensified over time (Fig. 4D, magenta arrowhead), indicating that some HMR-1 reaches the midline independent of PAR-3.

**Figure 4.**
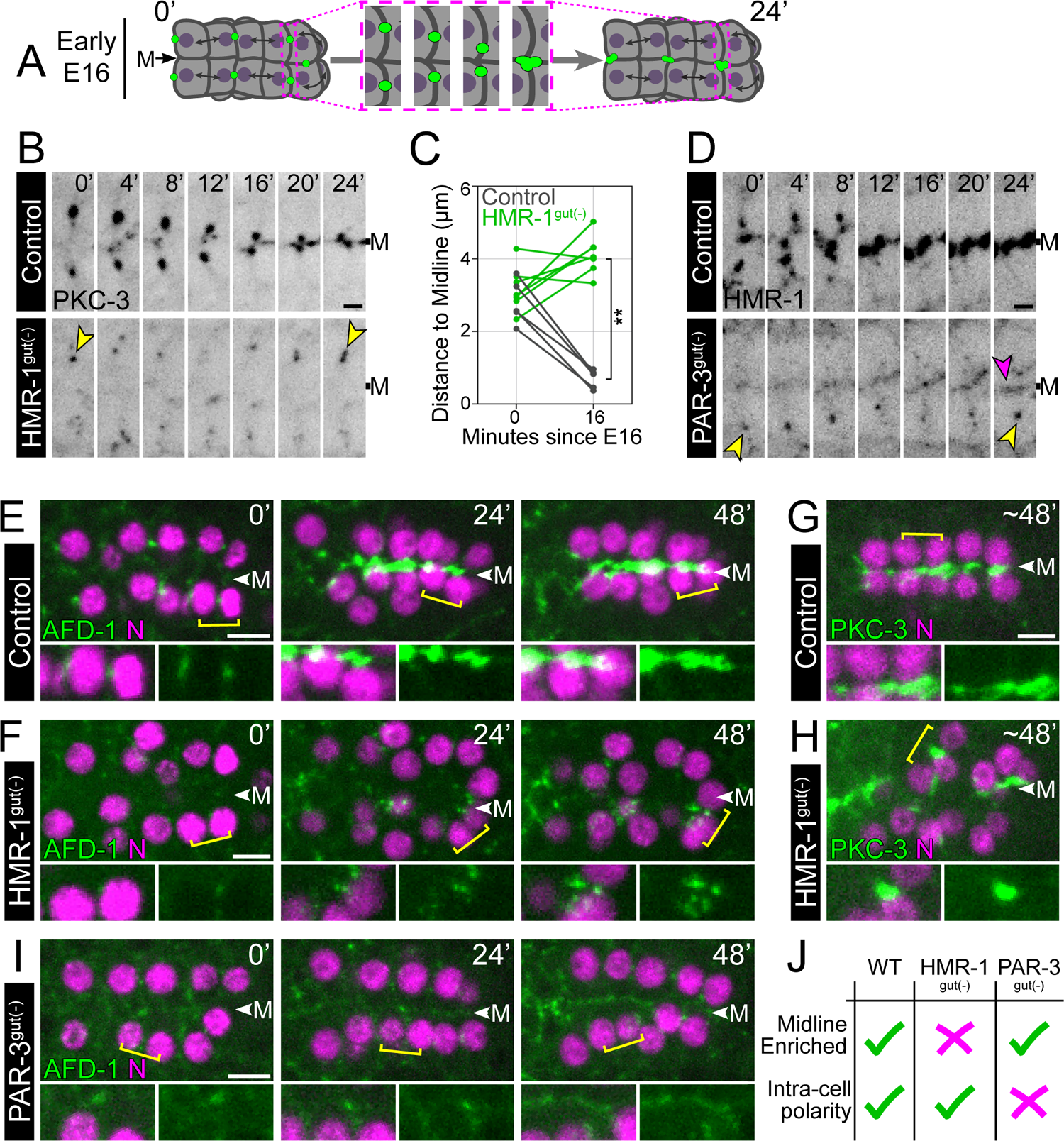
HMR-1 and PAR-3 accomplish tissue-level symmetry breaking through separable roles. A) Cartoon depicting dorsal views of LPC migration in E16 intestine with the two contacts shown in (A) and (C) indicated (dotted magenta box). B) Endogenous GFP:PKC-3 localization in Control (n=7) or HMR-1^gut(-)^ (n=9) embryos. 0’=First frame of E16. M = midline. Yellow arrowheads highlight LPC stalling. C) Analysis of PKC-3 migration over time. One dot pair represents one embryo. **p<0.01. D) Localization of endogenously GFP-tagged HMR-1 in Control (n=6) and PAR-3^gut(-)^ (n=8) embryos, labeled as in (A). Magenta arrowhead highlights HMR-1 enrichment at midline. E-I) Live dorsal views of E16 intestinal primordia. Yellow brackets indicate regions shown in insets. N = intestinal histone::mCherry. E-F, I) Endogenous AFD-1:GFP localization in Control (n=4), HMR-1^gut(-)^ (n=5), and PAR-3^gut(-)^ (n=8) embryos. 0’ = First frame of E16. G-H) Endogenous GFP:PKC-3 localization in Control (n=6) and HMR-1^gut(-)^ (n=12) embryos stage-matched to 48’ panels in E-F. J) Summary of PAR-3 and HMR-1 tissue-level functions. Scale bar=1 μm (B, D) or 5 μm (E-I; insets magnified 1.5x).

To further understand tissue-level polarity in these backgrounds, we visualized polarization over a longer time period using the adherens junction protein Afadin/AFD-1:GFP, the only LPC component found to cluster in both HMR-1^gut(-)^ and PAR-3^gut(-)^ animals. Like other LPC components, AFD-1 moved to the midline and spread across it shortly after the E8-E16 division in control embryos (Fig. 4E; Movie S2). Nuclei tracked closely with LPCs, due to dynein-directed movement along LPC-associated microtubules (Feldman and Priess, 2012). In HMR-1^gut(-)^ intestines, AFD-1 foci stalled at lateral membranes (Fig. 4F, similar to Fig. 4B).

Over time, foci from neighboring cells localized more closely and AFD-1 signal grew and intensified (Fig. 4F, insets; Movie S2). This mislocalized AFD-1 appeared to pull in associated nuclei, suggesting that cells aligned their polarity axes with immediate neighbors but not with respect to the midline (Fig. 4F, t=48 min). Indeed, large patches of PKC-3 accumulated at the center of these groups of nuclei, revealing that HMR-1^gut(-)^ intestines formed discontinuous clusters of apical islands in lieu of the continuous apical surface seen in controls (Fig. 4G-H; Movie S3). Importantly, the regions between these islands did not represent gaps between E cells, as cell membranes were always in contact (data not shown). Although apical islands persisted into E20, they tended to fuse and form a properly positioned yet variably discontinuous apical surface (see below). By the L1 larval stage, when control animals had a visible lumen, HMR-1^gut(-)^ animals instead had lumenal pockets separated by long lumen-free regions (Fig. S3A-A’). Thus, HMR-1^gut(-)^ animals possess dramatically mis-polarized intestines in which LPCs form and enlarge between neighbors but fail to migrate to the midline.

In contrast to HMR-1^gut(-)^ embryos, AFD-1 foci failed to enlarge in PAR-3^gut(-)^ embryos (Fig. 4I). Additionally, nuclei did not cluster toward AFD-1 foci but rather remained non-polarized within cells. Nevertheless, midline intensity of AFD-1 still increased through E16 with similar timing as control (Fig. 4I, insets; Movie S2). We note that some non-migratory AFD-1 foci still localized to lateral membranes during this midline enrichment, suggesting that distinct subpopulations of AFD-1 may exist with differing mobilities. By E20, midline-associated AFD-1 began assembling into junctional rings as previously reported (Pickett et al., 2021).

Despite their shared role in cellular symmetry breaking, these data indicate complementary but distinct roles for HMR-1 and PAR-3 in tissue-level symmetry breaking, consistent with their parallel roles in intestinal polarization at later stages of development (Pickett et al., 2021). PAR-3 is required to enlarge LPCs and connect them to other cellular elements such as the nucleus to facilitate cytoplasmic polarization while being dispensable for midline enrichment of junctional proteins. In contrast, HMR-1 is required to move LPCs to the midline but is dispensable for LPC growth (Fig. 4J).

### Tissue symmetry breaking requires the HMR-1 extracellular domain

We exploited the redundancy present in *C. elegans* adhesion to further investigate how HMR-1 could mediate polarity divorced from its adhesive functions. Generally, cadherins can fulfill signaling roles by clustering and interacting with neighboring cells through their extracellular domain while using an intracellular domain to interface with adaptors and the cytoskeleton. To investigate these potential roles, we overexpressed a truncated form of HMR-1 containing only its intracellular domain fused to a membrane tether in otherwise wildtype intestinal cells (ICD^PM^, Fig. 5A-A’). While ICD^PM^ was sufficient to mislocalize the HMR-1-binding protein HMP-2/β-catenin to all membranes (Fig. 5B), demonstrating functionality of the construct, ICD^PM^ did not mislocalize PAR-3 (Fig. 5C), suggesting that PAR-3 does not directly interact with HMR-1 or HMP-2. Upon depletion of endogenous HMR-1, ICD^PM^ also failed to polarize LPCs, as E8-stage embryos still lacked PKC-3 puncta and foci failed to migrate at E16 as in HMR-1^gut(-)^ embryos (Fig. 5D-F, 5G-I). Together, these data show that the HMR-1 intracellular domain is insufficient to drive LPC formation or migration, indicating that the extracellular domain of HMR-1 is required for its symmetry breaking functions.

**Figure 5.**
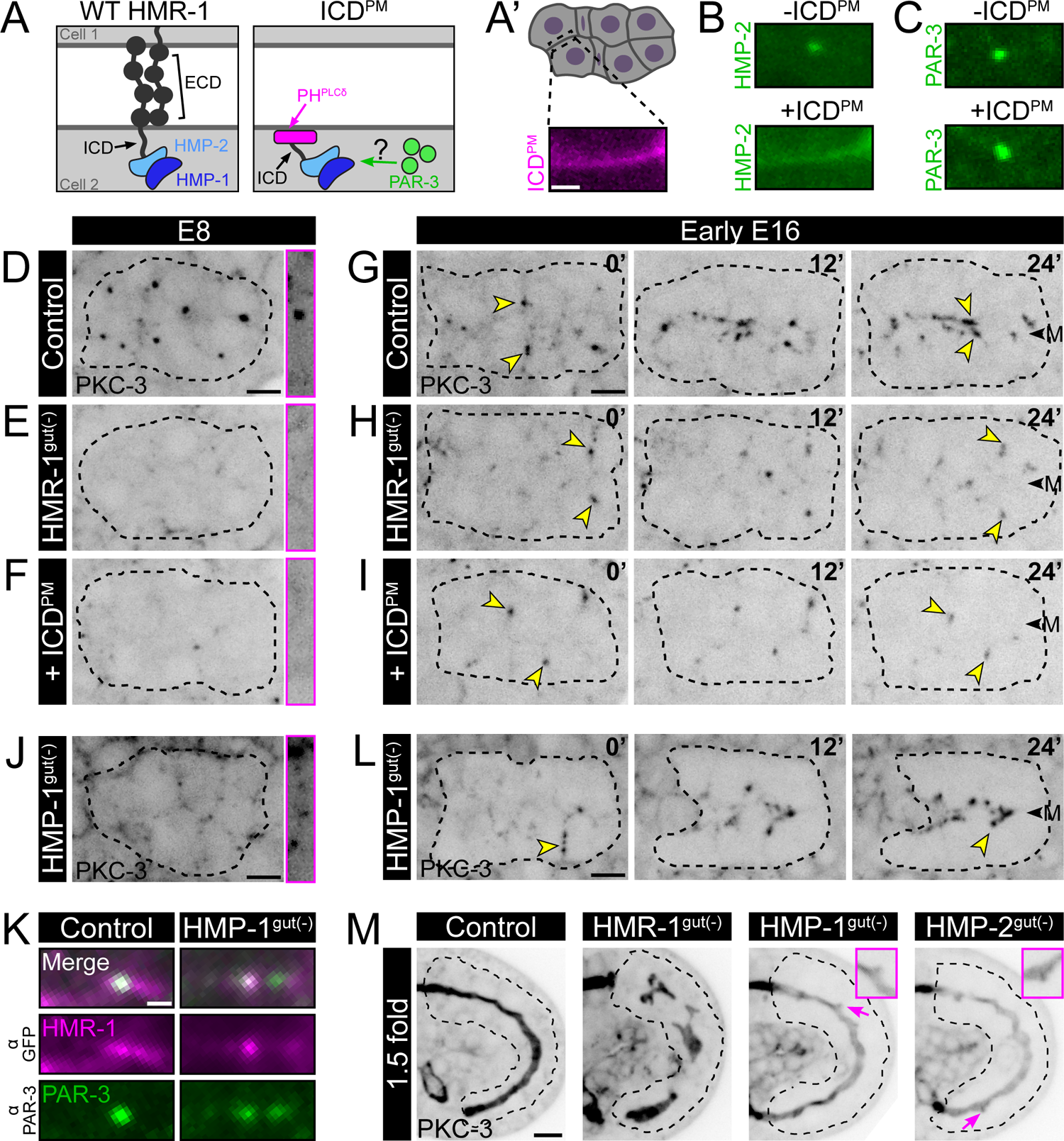
Symmetry breaking requires the HMR-1 extracellular domain but not the catenin adaptors HMP-2 or HMP-1. A) Cartoons of wildtype (left) and ‘ICD^PM^’ (right) HMR-1 constructs. ICD^PM^ binds the membrane via a PLC1δ1 plextrin homology domain (magenta) and is fused to mCherry. A’) Top: Cartoon of dorsal E8 intestine with indicated contact shown at bottom and in (B,C). Bottom: ICD^PM^ localization at one E8 contact in a live embryo (n>15). B-C) Live localization of HMP-2 or PAR-3 in control (HMP:2: n=14; PAR-3: n=4) or ICD^PM^-expressing (HMP:2: n=19; PAR-3: n=10) embryos. D-J, L) Live imaging of intestinal primordium (dorsal view; black dotted line) in embryos expressing endogenously GFP-tagged PKC-3. D-F, J) E8-stage Control (n=12), HMR-1^gut(-)^ (n=9), HMR-1^gut(-)^ + ICD^PM^ (n=5), or HMP-1^gut(-)^ (n=10) embryos. Magenta insets show one homotypic contact at high contrast. G-I, L) Early E16 stage in Control (n=5), HMR-1^gut(-)^ (n=5), HMR-1^gut(-)^ + ICD^PM^ (n=5), or HMP-1^gut(-)^ (n=7) embryos. 0’=First frame of E16. Yellow arrowheads track a subset of LPCs. ‘M’ = midline. K) Immunostaining for PAR-3 (α-PAR-3) and endogenously GFP-tagged HMR-1 (α-GFP) in Control (n>10) or HMP-1^gut(-)^ (n=7) E8 embryos, cropped to one homotypic contact as in A’. M) Live imaging side view of 1.5 fold stage Control (n=6), HMR-1^gut(-)^ (n=11), HMP-1^gut(-)^ (n=16), or HMP-2^gut(-)^ (n=15) embryos. Magenta arrows, insets highlight gnarled but continuous apical surfaces. Scale bars=2 μm (A’-C, K), 5 μm (D-L, M), or 1 μm (D-F). All insets magnified 1.5x.

### α- and β-catenin play a supportive but dispensable role in cellular symmetry breaking

HMR-1 interacts with the cytoskeleton and other effectors through HMP-2/β-catenin and HMP-1/α-catenin (Fig. 5A; Choi et al., 2015; Kang et al., 2017; Kwiatkowski et al., 2010), which comprise the adherens junction in mature epithelia. To determine whether symmetry breaking involves the full junctional complex, we next depleted HMP-1 or HMP-2 (Fig. S2B-C, S2E-F). At E8, the number of PKC-3 puncta in HMP-2^gut(-)^ embryos was reduced to a similar extent as following HMR-1 depletion (Fig. S3B-B’’, S3E), although the PKC-3 foci remaining after HMP-2 depletion were on average over twice as bright and 75% larger than those in HMR-1^gut(-)^ embryos (p=0.011 and 0.061, respectively; Fig. S3F). In contrast, HMP-1 depletion yielded smaller but more numerous PKC-3 foci and additional non-punctate membrane signal (Fig. 5D-E vs. 5J; Fig. S3E-F). Foci also contained HMR-1 and PAR-3 (Fig. 5K, Fig. S3C), suggesting that these foci are simply small LPCs. Thus, by E16 PKC-3 puncta were either fewer but brighter (HMP-2^gut(-)^) or smaller and more diffuse (HMP-1^gut(-)^) than in HMR-1 depletion. These distinct phenotypes are intriguing given that HMP-2 and HMP-1 function within the same complex, however such differences may reflect additional protein-specific binding interactions and functions.

Despite these deviations from the wildtype LPC pattern, PKC-3 was still swept to the midline following either HMP-1 or HMP-2 depletion (Fig. 5L and data not shown). At later stages, HMP-1^gut(-)^ and HMP-2^gut(-)^ embryos possessed widened and dim apical surfaces correlating with expanded and twisted lumens in larvae, but surprisingly embryos from both genotypes lacked the severe apical discontinuities present in HMR-1^gut(-)^ animals (Fig. 5M, S3A-A’’’). HMP-1 and HMP-2 therefore help to localize and consolidate LPC material during cellular symmetry breaking but appear dispensable for tissue-level symmetry breaking. While these milder phenotypes could formally reflect residual undegraded HMP-1 or HMP-2 below our detection limit, our data are in line with previous examination of *hmp-1(0)* maternal germline clones, which produced embryos with apparently normal intestines despite lacking functional maternal and zygotic HMP-1 (Costa et al., 1998). Furthermore, HMP-1 depletion successfully abrogated intestinal junctions as visualized by HMR-1:GFP (Fig. S3D, D’), as HMR-1 was not enriched in a junctional belt but localized more generally to membranes. Together, these data support the conclusion that HMR-1 plays an adherens junction-independent role in symmetry breaking.

### Symmetry breaking is not explained by differential composition or levels of candidate adhesion proteins

The information used by the intestine to break symmetry at homotypic contacts cannot be explained by the functions of HMR-1 and PAR-3 alone, as non-intestinal neighbors also express these proteins, raising the question of what information the intestine uses to enrich LPCs at homotypic contacts. We first tested whether this information involved an ‘adhesion code’ (Hashimoto and Munro, 2019; Röper, 2012) between homotypic and heterotypic contacts, but after examining several candidates that designate polarity in other contexts (Table S1), we were unable to find a protein whose localization or co-depletion with HMR-1 suggested involvement in intestinal polarity (Fig. S4). However, as over 40 adhesion proteins exist in *C. elegans* (Cox et al., 2004), we cannot rule out that other proteins working alone or in combination may still aid in symmetry breaking. We next tested whether contacts may differ in relative amounts of E-cadherin on the membrane, since cells in other contexts can self-organize based on cadherin levels alone (Duguay et al., 2003; Foty and Steinberg, 2005; Steinberg and Takeichi, 1994). Indeed, the E4 intestine expresses 24% more HMR-1 than its neighbors (assessed with a transcriptional *hmr-1p*::GFP reporter, Fig. S5A-B) and zygotic expression begins concomitant with the appearance of HMR-1 foci (Fig. S5C-D, arrowheads), leading us to hypothesize that elevated intestinal HMR-1 levels could kickstart LPC clustering preferentially at homotypic contacts. However, upon reducing intestinal HMR-1 we found only a mild phenotype: in HMR-1-depleted embryos retaining some non-degradable HMR-1 (‘HMR-1^gut(low)^’, Fig. S5E), LPCs were occasionally dim and slow to migrate (Fig. S5F-F’) but most assembled and migrated on time yielding normal polarity. Thus, while elevated intestinal HMR-1 levels may aid in reproducible LPC assembly, neither a difference in HMR-1 levels nor any of the tested adhesion molecules can explain why LPCs localize to homotypic contacts.

### LPCs are patterned by contact age and cell division

Insight into what makes homotypic contacts special in this primordium came from further characterization into the dynamics and behaviors of the LPC as the intestine underwent morphogenesis. As discussed above, LPCs form over a distinct timeline with HMR-1 and PAR-3 requiring 12-16 minutes and PKC-3 requiring 30 minutes to localize to LPCs (Fig. 2H-I). This timing suggests that cell/cell contacts must be stably associated for at least 30 minutes to establish a mature LPC. Although this gradual timeline was a hallmark of interphase LPC assembly, we observed more rapid assembly and disassembly behaviors during mitosis, once ventral E8 cells had dropped below their dorsal neighbors (Fig. 6A-A’). As E8 cells divided, two types of homotypic contact existed within the primordium: 1) those ‘on-axis’ with respect to anterior/posterior division orientation, closest to spindle poles (Fig. 6A’-B, blue), and 2) those ‘off-axis’ with respect to division, parallel to spindle poles (Fig. 6A’-B, yellow). LPCs at off-axis contacts rapidly dissipated during cell division (Fig. 6C, 6C’), while all of the LPCs that perdured and migrated at E16 (Fig. 1I) were at on-axis contacts (Fig. 6D, 6D’). Moreover, on-axis contacts appeared to receive LPC material specifically from divisions: at new homotypic contacts formed during the ventral descent of two E8 cells (Fig. S1A; Fig. 6A, white asterisks), LPCs were initially absent but arrived over about 8 minutes as cells completed anaphase (Fig. 6E, 6E’). Thus, while contact lifetime is a major contributor to gradual LPC formation in interphase, LPCs can also be quickly generated or destroyed by oriented cell division, affecting when and where LPCs form.

**Figure 6.**
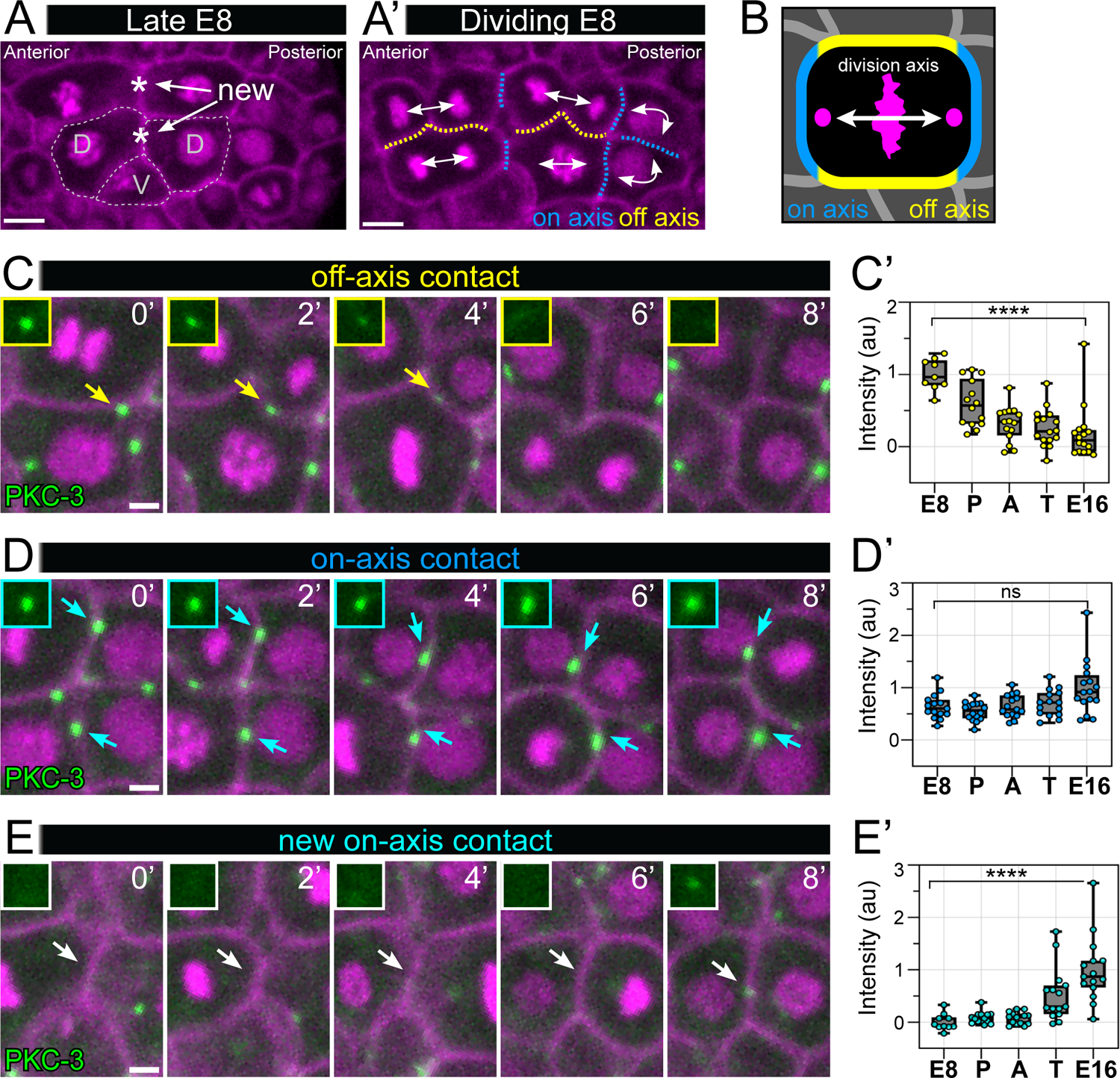
LPCs are patterned by oriented cell division. A) Live dorsal view of E8-stage intestinal primordium before synchronized divisions with histone::mCherry, membrane::mCherry. Ventral cells (‘V’) have dropped below dorsal neighbors (‘D’), creating new homotypic contacts (white asterisks). A’-B) Anterior/posterior division orientation creates ‘on-axis’ (blue) and ‘off-axis’ (yellow) contacts. The midline contact between posterior-most E8 cells is also on-axis due to oblique dorsal/ventral division of these cells (Sulston et al., 1983). C-E) Live timelapse imaging of off-axis, on-axis, and new on-axis homotypic contacts. Arrows point to disappearance (yellow), retention (blue), or appearance (cyan) of LPCs and are highlighted by insets. C’-E’) PKC-3 intensity measured during mitosis: P=prophase; A=anaphase; T=telophase. One dot represents one contact (n=8 embryos). ****p<0.0001. Scale bars=5μm (A-B) or 2 μm (C-E including insets).

### Contact duration and division orientation are sufficient to explain homotypic LPC enrichment

LPC dynamics observed at homotypic contacts in the intestine suggest a general set of rules for LPC formation and perdurance: 1) LPC maturation in interphase cells takes ∼30 minutes and therefore requires contacts of this age or greater to form; 2) LPCs at contacts on-axis with the spindle are reinforced; 3) LPCs at contacts off-axis with the spindle are erased. We therefore asked whether the application of these rules at heterotypic contacts could explain why we do not see LPCs at these surfaces. Intriguingly, we found that ephemeral LPCs did form at heterotypic contacts but were removed in two distinct ways. First, heterotypic puncta were removed as E/non-E contacts were broken and exchanged for contacts with new neighbors (Fig. 7A). Second, puncta were erased from heterotypic contacts as neighbors underwent division, most of which are oriented anterior/posterior and thus are off-axis with respect to these contacts (Fig. 7B).

**Figure 7.**
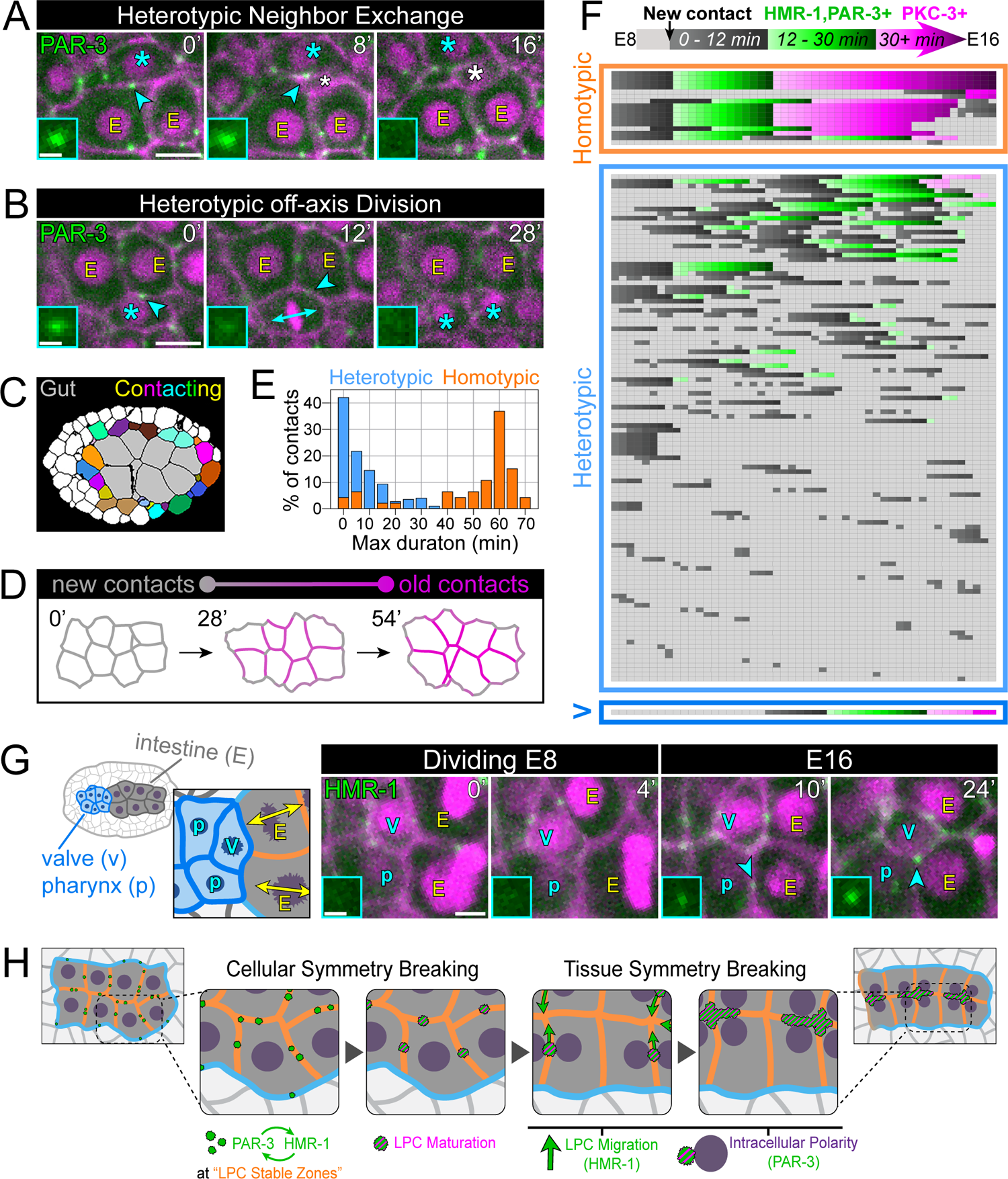
Contact duration and division orientation are sufficient to explain homotypic LPC enrichment. A-B) Dorsal view timelapse of endogenously GFP-tagged PAR-3 in an area at the interface between intestinal cells (‘E’) and a non-intestinal cell neighbor (cyan asterisk), with histone::mCherry, membrane::mCherry (magenta). Cyan arrowhead and inset highlight a heterotypic PAR-3 punctum that is removed when: A) the heterotypic contact breaks as a second non-E cell (white asterisk) intercalates dorsally or B) the heterotypic contact becomes off-axis as the non-E cell divides (cyan double arrow). C) Example E8-stage dorsal view segmented embryo used for analysis in D-F. D) Contact trace of intestinal membranes in an E8 stage primordium at three timepoints with relative contact age colored as shown. E) Percent of homotypic and heterotypic contacts from segmented embryos (n=3) binned by their maximum duration time, calculated from individually traced contacts. F) Heatmap of changing contact patterns in one segmented E8 embryo. Rows represent all points of cell/cell contact observed between E8 and E16, and columns represent 1.5-minute time steps within this period. Colors represent the occurrence and age of a contact (see key at top). The formation of a new contact at a specific time point is denoted by dark grey. Graded color changes indicate aging of each contact or on-axis division, while light grey indicates that the contact has been broken or has experienced an off-axis division. Green and magenta color shadings denote contact times long enough to allow localization of HMR-1/PAR-3 or PKC-3, respectively, based on our measured LPC assembly timeline (Fig. 2H). Shadings were accelerated following on-axis division to signify accelerated LPC recruitment. V = heterotypic valve/intestine contacts. G) Left: cartoon of dorsal view embryo indicating the interface between the E8 intestine and neighboring pharynx (p) and valve (v) primordia. Right: Timelapse of HMR-1 at this interface. Nuclei, membranes (magenta) as in A-B. H) Model of epithelial symmetry breaking. Scale bars=5 μm (A-B), 2 μm (G), or 1 μm (A, B, G insets).

Given the apparent effect of contact age and division axis on LPC perdurance, we used lattice light sheet movies of embryonic development (Cao et al., 2020) to track the movement and division of all cells surrounding the E8 intestine visible within a single plane (Fig. 7C; Movie S4). Homotypic intestinal contacts were stably associated, with most lifetimes >40 minutes (Fig. 7D-E). In contrast, heterotypic contacts experienced off-axis divisions and shorter lifetimes due to frequent neighbor exchanges (Fig. 7D-F; Movie S4): 94% of heterotypic contacts had a maximum duration less than 30 minutes, compared to just 16% of homotypic contacts with this contact duration (Fig. 7E). Within a given E8 embryo, only homotypic contacts perdured for longer than 30 minutes, the critical window required to localize HMR-1, PAR-3, and PKC-3 to LPCs in interphase intestinal cells (Fig. 7F). Intriguingly, we observed one notable exception to this pattern at the heterotypic contacts that form between anterior intestinal cells and the anterior pharyngeal valve cells, which lasted more than 30 minutes (Fig. 7F, ‘V’ panel; 7G, cartoon). Consistently, we observed LPCs at these heterotypic contacts at the end of E8 (Fig. 7G, t=0’, white arrowhead), which appeared to be further reinforced by their localization on-axis relative to the division of both the valve cell and the anterior intestinal cells (Fig. 7G). This contact surface is the basis for the connection that will form between the intestine and the pharynx, thus perduring LPCs at this site could help create a continuous digestive tract. Together, these data validate a model where asymmetry within a developing primordium is provided by contact lifetime and oriented division, information that is read out by the perdurance of LPCs at homotypic contacts.

## Discussion

Multicellular patterns are essential for organ function and animal development, yet the origins of many of these patterns are largely unknown *in vivo.* In this work, we identified a contact duration-dependent mechanism governing apico-basolateral polarization in the *C. elegans* intestinal epithelium. Our data support a model of symmetry breaking (Fig. 7H) informed by (1) differences in lifetime between homotypic and heterotypic contacts and (2) a molecular timeline of LPC maturation, which we propose is set by the intrinsic self-associating properties of the transmembrane protein HMR-1 and the scaffolding polarity protein PAR-3. In combination, these features allow for association of LPCs with long-lasting homotypic contacts, which act as ‘LPC stable zones’ to initiate cell- and tissue-level asymmetry. Within the LPC, HMR-1 and PAR-3 first act together to recruit apical cargo before playing separate roles in LPC migration and intracellular polarization, respectively, ensuring timely formation of a continuous apical surface across the primordium.

### The emergence of cell-level polarity through self-organizing components

We identified a stereotyped maturation sequence of the LPC as PAR-3 and HMR-1 initially appeared at E4 membranes as ephemeral foci before growing into brighter puncta that eventually localized the other apical determinants PAR-6 and PKC-3. Clustering is common among polarity proteins and reflects the general tendency of such molecules to self-associate (reviewed in Harris, 2017). For example, bud polarization in *S. cerevisiae* relies on Cdc42 clusters which form as slow-diffusing Cdc42 at the membrane interacts with fast-diffusing cytoplasmic Cdc42 (Chiou et al., 2017). PAR-3 and E-cadherin can similarly self-organize into nano- and micron-scale clusters (Dickinson et al., 2017; Harris, 2017; Pontani et al., 2016; Thompson et al., 2020; Troyanovsky et al., 2007; Yap et al., 2015), and we saw evidence that the self-organizing properties of these molecules underlies LPC assembly in the developing intestine: depleting either PAR-3 or HMR-1 prevented the other from growing into large puncta, but each protein on its own still localized to small foci like those observed at E4. Our results are consistent with data reported in the *Drosophila* blastoderm, where Par3/Baz interfaces with immature ‘spot’ junctions of E-cad/Shot (Harris and Peifer, 2004) upstream of aPKC (Harris and Peifer, 2005). Thus, our data support a conserved framework in which epithelial polarity is initiated and reinforced by self-organizing molecular principles.

### The LPC as a potential Apical Membrane Initiation Site?

While epithelia polarize using the same concert of proteins, these players are deployed in different ways depending on developmental context (Pickett et al., 2019). The symmetry breaking module we report here is similar in its composition and genetic interactions to polarizing spot junctions of the *Drosophila* blastoderm, but we observed clear differences in the number and stereotyped behavior of spot junctions and the LPC. New lateral membranes in the cellularizing blastoderm localize multiple spot junctions of variable size, which generally sweep toward the contact-free future apical surface (Harris and Peifer, 2005). In contrast, once associated with PKC-3, the LPC often appeared as a single perduring structure before undergoing directed movement toward the intestinal midline. LPC positioning along contacts was also not random but occurred midway along the length of each contact. In these ways, the LPC behaves less like a spot junction and more like the Apical Membrane Initiation Site (AMIS), an analogous singular structure comprised of apical and tight junction components seen prior to lumenogenesis (Bryant et al., 2010; Chen and Johnston, 2021; Rathbun et al., 2020). The AMIS is often placed at midbody remnants following mitosis, yet LPCs localize to both sister and non-sister homotypic contacts and are distinct from midbodies (data not shown). However, recent work demonstrated that cell divisions are not necessary for AMIS localization per se (Chen and Johnston, 2021; Liang et al., 2022) as a new, single AMIS still formed about halfway along the length of cell/cell contacts when division was blocked in mouse embryonic stem cells (Liang et al., 2022), akin to the LPCs we observed at E8. Moreover, both LPC and AMIS positioning are critical for tissue-level polarity: LPCs in HMR-1^gut(-)^ embryos stalled in place but continued to enlarge and polarize intracellular structures, leading to the formation of apical islands within the tissue. Similarly, MDCK or Caco-2 cells that fail to orient midbodies toward the cyst center have discontinuous apical surfaces and multi-lumen phenotypes (Jaffe et al., 2008; Lujan et al., 2017). However, while the AMIS is often positioned directly at the future apical surface, the LPC differs as it is born elsewhere and later moved to its correct location.

Given these similarities and differences between the LPC, spot junction, and AMIS, we propose that these structures are all variations on a conserved symmetry breaking machinery, with differences in components and polarization mechanisms dependent on context such as tissue anatomy and orientation of primordial cell divisions. For example, MDCK cells and Kupfer’s vesicle cells in zebrafish orient their divisions and cytokinesis toward their apical or future apical surface (Jaffe et al., 2008; Rathbun et al., 2020). In contrast, *C. elegans* intestinal cells, like most cells of the embryo at this stage, divide along the anterior-posterior axis in parallel to the future apical surface, thus we speculate that a mechanism based on assembly of an AMIS-like structure at all contacts followed by their directed migration may be the simplest solution to polarize. In support of this view, the mouse kidney nephron represents a similar context where polarization occurs in an aggregate of cells not undergoing division across a future apical surface, and like *C. elegans*, polarity emerges from the reorientation of membrane-associated PAR-3 patches toward the center of the primordium to create a lumen (Yang et al., 2013).

### E-cadherin as a transmembrane signaling platform

We report a symmetry breaking mechanism reliant on parallel inputs from PAR-3 and HMR-1, with PAR-3 elaborating polarity within individual cells and HMR-1 aligning polarity axes across cells. These proteins confer separable functions to a single module, a conceptually straightforward and likely conserved (Bulgakova et al., 2013; Houssin et al., 2021; Valdivia et al., 2020) mechanism to achieve cell- and tissue-level polarity. E-cadherin localization to cell/cell contacts has previously been proposed to initiate epithelial polarity (Nejsum and Nelson, 2007, 2009), but specific requirements for E-cadherin in symmetry breaking have been difficult to divorce from its often-essential adhesive function. However, the roles for HMR-1 described here occur separately from both adhesion and the mature adherens junction while still requiring the extracellular domain, thus we propose a more general early function for HMR-1 as an intercellular signaling platform which orients intracellular polarity machinery toward a common tissue surface.

How do HMR-1 platforms assemble and interface with PAR-3? E-cadherin is known to cluster through its extracellular domain, which not only strengthens adhesions (Harrison et al., 2011; Pontani et al., 2016; Thompson et al., 2020) but also creates local signaling pools. For example, HMR-1 clusters in the one-cell *C. elegans* embryo locally downregulate myosin to preserve integrity of the actomyosin cortex (Padmanabhan et al., 2017). *In vitro,* E-cadherin can selectively segregate proteins from each other, concentrating them into membrane-associated corrals (Troyanovsky et al., 2021). HMR-1 may similarly package PAR-3, although any interaction between these proteins is likely indirect. While Par3/Baz can bind several adherens junction proteins in *Drosophila* (Harris and Peifer, 2005; Wei et al., 2005), such interactions have not been observed in *C. elegans*, and we found that overexpressing the HMR-1 intracellular domain failed to alter PAR-3 localization. Possibly, HMR-1 and PAR-3 may populate distinct subdomains in the LPC as they do at spot junctions (McGill et al., 2009), a model that will require future higher resolution imaging studies to test.

Surprisingly, this signaling function of HMR-1 was largely independent of its intracellular adaptors HMP-1/α- and HMP-2/β-catenin, which aided in LPC assembly but were not required for overall polarity. The catenins link E-cadherin to the actin cytoskeleton, and actin is also dispensable for intestinal polarization (Feldman and Priess, 2012). In other systems, actin can behave as a molecular fence to guide or limit E-cadherin clustering (Hong et al., 2013; Wu et al., 2015). As HMP-1 or HMP-2 depletion affected the size and regularity of LPCs, we speculate a similar mechanism may exist here to corral HMR-1 into puncta. Clustering may also be mediated by actin-based flow along membranes toward the center of each contact, similar to flows that incorporate E-cadherin into adherens junctions (Woichansky et al., 2016). Despite their small size, foci in HMP-1^gut(-)^ embryos were still swept to the midline on time, raising the question of why LPCs normally enlarge into puncta if smaller clusters can also polarize. We propose that larger clusters of HMR-1 and PAR-3 may more efficiently recruit factors necessary for maturation and migration, but that clustering arises through parallel mechanisms and is therefore perturbed but not eliminated in HMP-1^gut(-)^ embryos. Indeed, HMP-1^gut(-)^ foci still localized HMR-1, PAR-3, and PKC-3. Some minimal degree of clustering may be required for the transport of these proteins, as also observed in the *C. elegans* zygote (Chang and Dickinson, 2022), but perhaps further abrogating clustering is needed to uncover such defects. For example, the p120 catenin homolog JAC-1, which was not tested here, could act on its own or in parallel to HMP-1 and HMP-2 to further promote clustering and polarization.

As the intestine polarizes independent of α-, β-catenin and actin, HMR-1 may instead orchestrate LPC migration through microtubules. LPCs enlarge as a new microtubule organizing center is established at this site, and the stalled LPC phenotype seen in HMR-1 depletion resembles the migration delay reported upon microtubule perturbation (Feldman and Priess, 2012; Leung et al., 1999). HMR-1 could actively transport LPCs to the midline by linking to microtubules, or microtubules may deliver HMR-1 itself to homotypic contacts, as dynamic microtubules can enrich E-cadherin on membranes (Stehbens et al., 2006). HMR-1 could also act independent of microtubules, perhaps through a general midline-directed membrane flow. Even upon PAR-3 depletion, HMR-1 gradually enriches at the midline, consistent with a model where a flow-responsive pool of HMR-1 normally engages less-mobile components like the PAR proteins to enable their migration.

### Contact lifetime differences as a symmetry breaking cue – a temporal component to differential adhesion

The *C. elegans* intestine must break symmetry while fully surrounded by other cells and lacking cues from a basement membrane or midbody (Graham et al., 1997; Leung et al., 1999), raising the question of what asymmetric information this tissue uses to generate polarity axes. One emergent theme to polarize internal tissues is through cell-type specific expression of adhesion molecules that enrich homotypically and recruit effectors to pattern morphogenesis (Hashimoto and Munro, 2019; Klompstra et al., 2015; Röper, 2012; Sidor et al., 2020). However, we were unable to find a suitable candidate to delineate intestinal homotypic from heterotypic contacts. Instead, we demonstrated that the short-lived contact duration and anterior/posterior division orientation of most intestinal neighbors was sufficient to explain their lack of persistent LPCs, thus defining a contact asymmetry within the primordium.

What imparts extra stability to homotypic contacts? One explanation would involve a higher level of adhesion inside the intestine than outside of it, a ‘differential adhesion’ phenomenon responsible for tissue patterning in many contexts (Tsai et al., 2022). Consistent with this model, the NCAM ortholog NCAM-1 localized within the E8 primordium but not in neighbors, suggesting that homotypic contacts are potentially more adhesive. E cells also express 24% more HMR-1 on average than neighbors, a difference in the level of adhesion protein required for spontaneous sorting of cultured L cells (Duguay et al., 2003). However, adhesion differences cannot solely explain symmetry breaking, since we also observed heterotypic LPCs removed by active migrations, intercalations, and divisions. Intestinal neighbors give rise to various tissue types including body wall muscle, hypodermis, seam cell, and neuron. These precursors actively migrate within the embryo during development (Sulston et al., 1983), but the details of such migrations remain largely uncharacterized. Given our observations, however, we propose that homotypic/heterotypic contact stability differences are in part a byproduct of the inherently distinct differentiation programs executed by intestinal neighbors during morphogenesis.

The differences in homotypic/heterotypic contact stability that break symmetry may be a simple consequence of many different modes of development. Conceptually, groups of cells with similar developmental trajectories are likely to associate with one another for longer since they undergo programmed divisions and migrations at the same time and in response to the same cues. Such preferential associations of like cell types throughout development impart a ‘temporal’ component to differential adhesion, where homotypic and heterotypic differences in contact lifetime can be explained not only by adhesion molecule composition but also as breaks in contact necessarily occur when cells undergo their own morphogenesis programs. By coupling relative stability of homotypic contacts to the intrinsic assembly timeline of polarizing components, the *C. elegans* intestine reflects a tissue-intrinsic symmetry breaking paradigm. As all internally developing epithelial primordia express PAR-3 and E-cadherin and must polarize within a milieu of other cells, we believe this mechanism reflects a generalizable mode of animal tissue patterning.

## Supporting information

Combined SI

## Acknowledgements

We thank Jeremy Nance, Mike Boxem, Ken Kemphues, Dan Dickinson, Michael Nonet, and Maria Sallee for worm strains and plasmids. We also thank members of the Feldman lab for helpful discussions about the research and manuscript. Some strains were provided by CGC, funded by the NIH Office of Research Infrastructure Programs (P40 OD010440). This work was funded by CMB Training Grant T32 GM007276 and a Stanford Graduate Fellowship to VFN, F32 GM129900-01 awarded to MAP, and an NIH New Innovator Award DP2 GM119136-01 and RO1 GM133950 awarded to JLF.

## Author Contributions

Conceptualization, J.L.F. and V.F.N.; Methodology, V.F.N., M.A.P., J.L.F.; Formal Analysis, V.F.N.; Investigation, V.F.N., J.L.F., M.A.P.; Writing – Original Draft, V.F.N. and J.L.F.; Writing – Review & Editing, V.F.N., J.L.F., M.A.P.; Visualization, V.F.N. and J.L.F.; Supervision, J.L.F.; Funding Acquisition, J.L.F., V.F.N., M.A.P.

## STAR Methods

### *C. elegans* strains and maintenance

Nematode strains were cultured on Nematode Growth Medium (NGM) plates coated with a lawn of *E. coli* OP50 (Sulston and Brenner, 1974) unless otherwise stated. For all experiments, animals were maintained at 20°C. Strains used in this study are listed in the Key Resources Table.

### CRISPR/Cas9 cloning and editing

CRISPR alleles were generated using the Self-Excising Cassette (SEC) method (Dickinson et al., 2015). Plasmid pDD162 was used to deliver Cas9 and a specific sgRNA, inserted into the plasmid with the Q5 Site-Directed Mutagenesis Kit (NEB). To create homology directed repair templates, 5’ and 3’ homology arm sequences were amplified using Phusion High-Fidelity DNA Polymerase (Thermo Scientific), and these were cloned into the appropriate SEC vector (pFJ250 for ZF:GFP tag; pDD282 for GFP tag) using Gibson Assembly (NEBuilder HiFi DNA Assembly Master Mix, NEB). Note that GFP or ZF:GFP alleles also contain 3xFLAG. For each edit, a mixture of the repair template (25 ng/μl), sgRNA/Cas9 plasmid (50 ng/μl), and pBlueScript (57.5 ng/μl) were injected into young adult *zif-1(gk117)* hermaphrodites. New CRISPR edits were recovered from injected worms as previously described (Dickinson et al., 2015) and back-crossed twice prior to use in experiments. New CRISPR alleles generated during this study and their associated plasmids, sgRNA sequences, and homology arm primers are listed in Table S2.

### Transgene cloning and editing

All transgenic lines were generated by injecting plasmid constructs into young adult hermaphrodites. pVN68 was made by cloning the PH domain from PLC1delta1 and the HMR-1 intracellular domain corresponding to amino acids 1151-1269 (similar to a construct reported in Klompstra et al., 2015) into a backbone between the *end-1* promoter and mCherry. Extrachromosomal ‘ICD^PM^’ arrays *wowEx210, wowEx211, wowEx212* were made by injecting a mixture of 5 ng/μL pVN68, 50 ng/μL pMP1, and 50 ng/μL pRF4. pVN56 was produced by cloning the *hmr-1* promoter::5’ UTR, GFP, and the *hmr-1* 3’UTR into pLF3FShC. *wowIs30* was generated by injecting a mixture of 50 ng/μL pVN56 and 50 ng/μL pBS and obtaining a single-copy insertion using the RMCE integration method (Nonet, 2020). The extrachromosomal *hmr-1* ‘rescuing’ array (*wowEx126*, used in Fig. 3D) was generated by injecting a mixture of 2 ng/μL pJN455, 2.5 ng/μL pDD401, and pBS to 150 ng/μL. *wowEx113* was generated by injecting a mixture of 50 ng/μL pJF248, 50 ng/μL pSA109, 2.5 ng/μL pCFJ90, and 47.5 ng/μL pBS. *wowIs28* is a spontaneous integrant of *wowEx113* and has been used previously (Sanchez et al., 2021).

### Embryo staging considerations

The following morphological landmarks were used to ensure embryos within each experiment were similarly staged: (1) completion of E4-E8 division at 205-210 mpf; (2) ventral movements of Eara and Earp at ∼245-250 mpf (Asan et al., 2016; Leung et al., 1999); (3) E8-E16 division at 270-285 mpf. E8-stage LPC assembly defects (Fig. 3, Fig. 5D-F, J) were assessed between 250 mpf and 265 mpf. Movies of tissue-level symmetry breaking in control and depletion backgrounds (Fig. 1I, Fig. 4, Fig. 5G-I, Fig. 6) were started at ∼280 mpf.

### Microscopy

For all imaging, 1-2-day old adults were incubated in M9 solution for 2-4 hours at 20°C. Adults were cut open to release embryos which were subsequently used for live imaging or staining. For live imaging, embryos or L1 larvae were mounted on a pad made of 3% agarose dissolved in M9 solution. L1 animals were treated with 2 mM levamisole for 3-5 minutes prior to mounting. Both live imaging and fixed samples were imaged on a Nikon Ti-E inverted microscope (Nikon Instruments) with a confocal spinning disk head using a 60x PLAN APO oil objective (NA = 1.4) controlled using NIS Elements software (Nikon). Images were captured using an Andor Ixon Ultra back thinned EM-CCD camera controlled using NIS Elements software (Nikon) at a z-sampling rate of 0.5 μm and using 488 nm, 561 nm, or 405 nm imaging lasers. All images were processed in Fiji/ImageJ and Adobe InDesign or Photoshop. All imaging and processing parameters (laser power, exposure time, LUTs) were kept constant within experiments.

### Degradation

All depletion strains were maintained with *zf::gfp* alleles over the appropriate balancer (see strains list in Key Resources Table), with the exception of JLF735 where degradable HMR-1:ZF:GFP was supplemented with the rescuing array *wowEx126[hmr-1p::hmr-1::gfp]*. For all experiments, gut(-) animals were taken from 1-2 day old Balancer(-) or wowEx126(-) mothers to ensure both maternal and zygotic pools of each protein were ZF:GFP-tagged. We note that no available balancer covered the *hmp-2::zf::gfp* locus, so this allele was maintained over *tmC27.* To ensure HMP-2^gut(-)^ embryos contained no untagged HMP-2 resulting from recombination of this unbalanced allele, we verified that Balancer(-) adults were homozygous for *hmp-2* before imaging their embryos. We verified loss of intestinal fluorescence upon degradation of HMR-1:ZF:GFP, ZF:GFP:HMP-1, and HMP-2:ZF:GFP (Fig. S2D-F), and PAR-3:ZF:GFP intestinal depletion as has been characterized previously (Pickett et al., 2021). While ZIF-1-mediated depletion is not equivalent to a genetic null, we and others consistently see strong depletion of targets and expected depletion-associated phenotypes (this study; Abrams and Nance, 2021; Liang et al., 2020; Magescas et al., 2021; Pickett et al., 2021; Sallee et al., 2018, 2021; Sanchez et al., 2021). We note that our HMP-1-tagging strategy leaves undegraded one predicted isoform, although this isoform is likely nonfunctional as it lacks HMP-2 or actin binding domains and only encodes a predicted disordered region (Fig. S2C). Most depletions were performed with *wowEx113[elt-2p::ZIF-1]*. In some depletion backgrounds, Balancer(-); *wowEx113+* animals were too rare to efficiently image. In these cases, we achieved degradation using *wowIs28,* a spontaneous integrant of the *wowEx113* array, which degrades ZF:GFP-tagged proteins with similar efficiency (Fig. S2G). Most degradation experiments were paired with controls which expressed the ZIF-1 machinery (*wowEx113* or *wowIs28*) and relevant markers but lacked any *zf::gfp*-tagged alleles. Two exceptions to this were: (1) Fig. 3D-D’, where HMR-1^gut(-)^ embryos were paired with a control which retained *hmr-1::zf::gfp* but lacked *wowIs28*; (2) Fig. 4H-I and Fig. 5M, where HMR-1^gut(-)^ embryos were paired with a control expressing non-ZF:GFP-tagged HMR-1 provided by a balancer.

### Generating HMR-1^Z1/2^ or HMR-1^gut(low)^ animals

To generate HMR-1^Z1/2^ embryos (Fig. S5C-D), JLF503 males were crossed to *tmc27, wowEx113(-)* JLF811 hermaphrodites. To generate HMR-1^gut(low)^ embryos (Fig. S5E-F’), JLF906 males were crossed to *tmC27(-), wowEx113(-)* JLF812 hermaphrodites. After one or two days, hermaphrodites were incubated in M9 for 3-4 hours and their embryos were released for imaging. Embryos were confirmed as cross progeny by the presence of HMR-1:GFP signal in later-stages (for HMR-1^Z1/2^) or by the presence of *elt-2p:histone:mCherry* (for HMR-1^gut(low)^).

### Immunofluorescence

1-2-day old adults were incubated in M9 solution for 2-4 hours at 20°C. Adults were cut open to release embryos, which were mounted on poly-L-lysine-coated microscope slides containing Teflon spacers (Leung et al., 1999) and frozen on dry ice. Embryos were permeabilized with the freeze-crack method and fixed in methanol for 5 minutes at −20°C. Following 2 5-minute rinses with PBS and 1 5-minute rinse with PBT (PBS + 0.1% Tween), embryos were incubated in primary antibody overnight at 4°C. Embryos were then rinsed 3 times for 5 minutes each in PBT and incubated in secondary antibody for 1 hour at 37°C. Embryos were rinsed once in PBT, twice more in PBS, and then were mounted in Vectashield (Vector Laboratories) and stored in the dark at room temperature. The following primary antibodies were used: anti-PAR-3 (DSHB, 1:25); anti-GFP (Abcam, 1:200). The following secondary antibodies were used: Cy3-anti-mouse (Jackson Immunoresearch Laboratories, 1:200); 488-anti-rabbit (Jackson Immunoresearch Laboratories, 1:200). Nuclei were visualized with DAPI (Sigma, 1:10,000).

### RNAi treatment

RNAi was performed by feeding larvae with HT115 bacteria transformed with *LacZ(RNAi)* plasmid (Ahringer, 2006) or *pop-1(RNAi)* plasmid (generated in Rual et al., 2004). NGM plates supplemented with IPTG and Ampicillin (Ahringer, 2006) were seeded with HT115 cultures and grown 48-72 hours in the dark at room temperature. L3-L4 stage larvae were grown on these plates for 48-72 hours at 25°C, and their progeny were imaged live as described above.

### Quantification

#### General considerations

Only embryos with dorsal surfaces closest to the coverslip were used for quantification, and all LPC measurements were taken from only the dorsal tier of intestinal cells. Unless otherwise stated, each LPC was measured using the single Z slice which captured the most signal. All projections, ROIs, and measurements were performed in Fiji/ImageJ. Membrane:mCherry was used to accurately draw ROIs unless otherwise stated.

#### Counting number of LPCs per contact

Maximum intensity measurements were taken in the cytoplasm (background) and at every region along homotypic and heterotypic contacts where PKC-3 signal appeared higher than background. Each of these regions was classified as an LPC if its intensity was greater than 5 standard deviations above average background signal (an empirically determined threshold, which consistently categorized regions as an LPC or not in agreement with visual observations). Contacts were then categorized as having 0, 1, or 2 LPCs based on this threshold (Fig. 1F).

#### Quantifying LPC diameter, position along contact, and position relative to membranes

To calculate LPC diameter (Fig. 1G) and position along contact (Fig. S1C), a line scan was generated from an ROI drawn along each contact through an LPC (marked by PKC-3). From each curve, (1) LPC diameter was approximated by calculating the Full Width at Tenth Max; (2) LPC position along contact was measured by recording the distance (x) corresponding to maximum PKC-3 intensity (y) and converting this into a relative position from 0-100%. To measure the position of PKC-3 relative to membranes (Fig. 1H), a second line scan was generated from an ROI drawn perpendicular to each contact through an LPC. Both PKC-3 and membrane intensity were measured, and all recorded line scans were aligned such that peak membrane signal was set to x=0.

#### Measuring LPC intensity at new homotypic contacts or during mitosis

To characterize LPC genesis, PAR-3, HMR-1, or PKC-3 punctum intensity was measured at new E8-stage contacts (Fig. 2H) with t=0’ defined as the first time point when new membrane was visible following E4-E8 divisions (see Fig. 2E cartoon). To assess LPCs during E8-E16 cell division, PKC-3 punctum intensity was measured at: E8 prior to division, prophase, anaphase, telophase, and E16 just following division. Staging was approximate, as contacting E cells rarely divided in perfect synchrony. For both quantifications, ROIs of constant size were drawn at each contact to measure average intensity (I) of (1) the brightest observable punctate signal, I^LPC^, (2) an area on the contact clearly devoid of puncta, I^Membrane^, (3) the edge of the slide, I^bkgd^. Punctum Intensity was calculated as the ratio of (I^LPC^ – I^bkgd^) / (I^Membrane^ – I^bkgd^) – 1, and these values were averaged across contacts and embryos. To plot LPC intensity changes during division, values were normalized such that the average E8 signal (Fig. C’) or the average E16 signal (Fig. D’, E’) was set to 1.

#### Average PKC-3 intensity in control and depletion backgrounds

A line ROI was drawn along the length of each homotypic contact (excluding corners and vertices) and a line scan was taken of PKC-3 intensity. Two more ROIs were drawn to measure average intensity at (1) a region on the membrane clearly devoid of LPCs, I^Membrane^, and (2) the edge of the slide, I^bkgd^. I^puncta^ was calculated by summing along the line scan any pixel > I^Membrane^. PKC-3 intensity was calculated as the ratio of (I^puncta^ – I^bkgd^) / (I^Membrane^ – I^bkgd^) – 1, and these values were averaged within embryos (Fig. 3B).

#### PKC-3 puncta number, size, and intensity in control and depletion backgrounds

For all images, 6-slice summed projections were taken that best captured dorsal E8 nuclei. In each embryo, a polygon ROI was drawn to encompass all dorsal nuclei. An intensity threshold was then applied to PKC-3 signal within the ROI. The threshold was calculated as the average intensity value corresponding to the top 1.5% brightest PKC-3 signal in control embryos (empirically determined to capture the most real LPCs while minimizing noise). After applying the threshold across all control and depletion backgrounds, the ‘Analyze Particles’ function (Fiji/ImageJ) was used to identify a punctum as any particle > 3 pixels with circularity > 0.5. This function output the number of PKC-3+ puncta per embryo (Fig. S3E), as well as the intensity and area of each individual punctum (Fig. S3F)

#### PKC-3 distance to midline during tissue-level polarization

A line was drawn from the brightest PKC-3 punctum on the membrane to the closest point on the midline. The length of this line was reported as Distance to the Midline in μm (Fig. 4C). Measurements were taken starting at the first time point following completion of E8 cell divisions (t=0’). Only posterior-most LPCs were used for this analysis (as in Fig. 4A-B).

#### hmr-1 transcriptional levels ratio

Summed *hmr-1p*:GFP intensity was measured in (1) an ROI drawn around the E4 intestine (I^intestine^) and (2) an ROI drawn around immediate intestinal neighbors (I^intestine+neighbors^). I^neighbors^ was calculated by subtracting I^intestine^ from I^intestine+neighbors^. All summed *hmr-*1p:GFP intensities were converted to average intensities by dividing by the area of each region, and the ratio of Average I^intestine^/Average I^neighbors^ was taken.

#### Comparing degradation efficiencies of wowEx113 and wowIs28

7-slice summed projections were made surrounding a pre-defined ‘midpoint’ Z slice where dorsal E nuclei were brightest. In each embryo, an intestinal ROI was drawn as a freeform shape which encompassed all visible E nuclei. Nuclei were easily seen in *elt-2p::zif-1* strains, which also expressed intestinal histone:mCherry, and in control embryos nuclei could be visualized by boosting LUTs to reveal nuclei as ‘black holes’ devoid of any GFP. Average GFP intensity was measured both inside this ROI and elsewhere on the slide to account for background, and similar measurements were made in JLF155 animals to estimate intestinal autofluorescence. Intestinal GFP intensity was approximated by subtracting slide background and autofluorescence from the average intestinal ROI intensity (Sallee et al., 2018).

#### Contact duration

Cell/cell ‘contact duration’ measurements here refer to the amount of time a contact existed without being broken and without either cell undergoing an off-axis division. Analyses were performed using previously published membrane-segmented lattice light sheet movies of *C. elegans* embryonic development (Cao et al., 2020). Movies had 1.5-minute timesteps and were cropped to start after E4-E8 divisions and end after E8-E16 divisions. For each time point, a single Z slice was chosen which approximately captured a middle section through the dorsal tier of the intestine. All cells contacting the intestine were tracked for the duration of E8, using both the single Z-slice movie as well as the raw multi-Z step data, and each of these cells was manually labeled a unique color in Microsoft Paint (Fig. 7C, Movie S4). Only non-mitotic cells in contact with the intestine were labeled, thus many cells lost and regained their color label through the movie as they made and broke contact or entered mitosis. Python script *contact_durations.py* was next used to recognize and isolate unique color labels in the movie, so that individual contact durations could easily be counted. For each labeled cell, contact durations were measured by counting the number of time steps for which that cell remained in contact with a specific intestinal cell. Of these values, the longest contact time was recorded as the Maximum Contact Duration (plotted in Fig. 7E).

#### Contact tracing and heatmap construction

For one representative embryo, individual contact duration values were visually depicted as a ‘Contact Trace’ (Fig. 7D, Movie S4): movie stills were imported into Adobe Illustrator, each contact was manually traced, and each tracing was color-coded from grey to magenta corresponding to its duration. This data was additionally incorporated into the heatmap shown in Fig. 7F, where rows correspond to the number of unique contacts to exist at any time during the contact trace movie, columns represent 1.5-minute time steps within the movie, and shading represents the presence of a specific contact at a given time. Using our previous timing analysis of LPC assembly (Fig. 2H), we overlaid onto this heat map our expectations about whether each contact had existed long enough to accumulate a mature LPC (PAR-3+, HMR-1+, PKC-3+; contact > 30 minutes, shaded in magenta), an immature LPC (PAR-3+, HMR-1+, PKC-3–; contact between 12 and 30 minutes, shaded in green), or no LPC components (PAR-3–, HMR-1–, PKC-3–; contact < 12 minutes, shaded in grey). For contacts experiencing on-axis divisions, the time point following this division was given a magenta shading to signify accelerated LPC assembly. Heatmap was constructed manually using Adobe Illustrator.

#### Statistical Analyses

GraphPad Prism 9.3.1 was used for all graphs and statistical tests. We performed most analyses using the Kruskal-Wallis test, a non-parametric alternative to the one-way ANOVA test, followed by the post-hoc Dunn test for multiple comparisons. For comparisons between only two samples, the Mann Whitney U Test was performed. We used the one-sample Wilcoxon test to determine whether the ratio of intestinal to non-intestinal *hmr-1p:*GFP expression significantly differed from 1 (Fig. S5B).

## Supplemental Movies

**Movie S1. LPCs perdure through divisions and seed the future apical surface.** Cropped dorsal view timelapse of 4 E8 cells. ‘M’ designates the tissue midline, and yellow arrowheads highlight LPC retention and migration. Scale bar = 5 μm.

**Movie S2.** HMR-1 and PAR-3 have separable roles in LPC migration and intracellular polarization. Endogenous AFD-1:GFP localization in a Control, HMR-1^gut(-)^, and PAR-3^gut(-)^ embryo. 0’ = First frame of E16. ‘M’ designates the tissue midline. Scale bar = 5 μm.

**Movie S3.** PKC-3 forms apical islands following HMR-1 depletion Endogenous GFP:PKC-3 localization in a Control and HMR-1^gut(-)^ embryo. 0’ = First frame of E16. ‘M’ designates the tissue midline. Scale bar = 5 μm.

**Movie S4.** Homotypic contacts are longer-lasting than heterotypic contacts. Left: Timelapse of E8-stage embryo with membranes segmented (Cao et al., 2020). Intestinal cells are depicted in dark grey, and all cells in contact with the intestine are also colored. Colors are lost from cells that lose contact with the intestine. Cells undergoing mitosis are shaded in light grey. Right: A tracing of all E8 cell membranes. Membranes incrementally change color from grey to magenta with every consecutive time point that they are in contact with the same neighboring cell. When contact is broken or at least one neighbor undergoes mitosis, the membrane tracing is reverted to the starting grey color denoting a new contact.

