## Supplementary material for "E-cadherin/HMR-1 and PAR-3 break symmetry at stable cell contacts in a developing epithelium": Combined SI

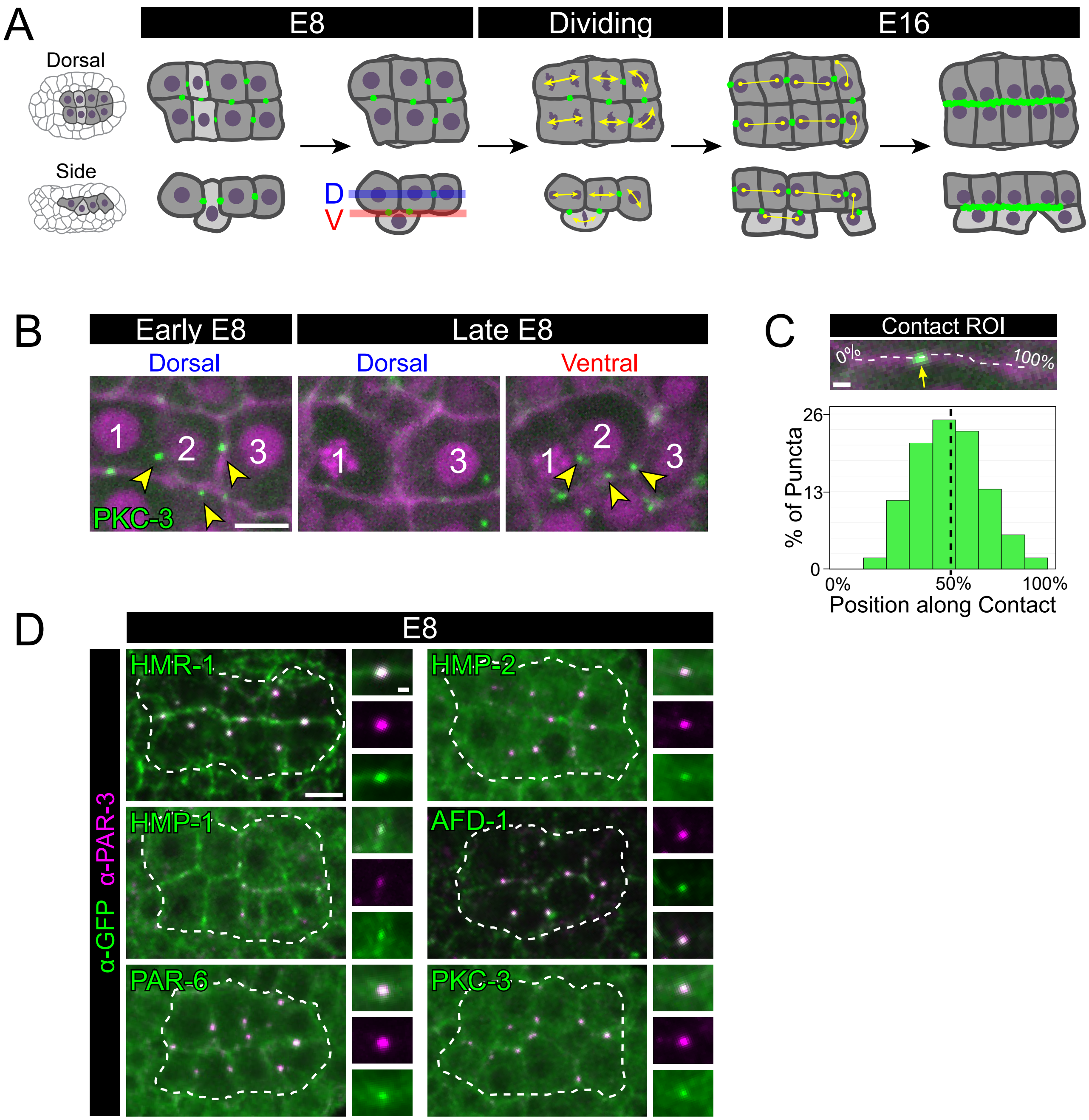
A
E8
Dividing
E16
Dorsal
Side
D
V
B
Early E8
Late E8
Dorsal
Dorsal
Ventral
1
2
3
1
3
1
2
3
PKC-3
C
Contact ROI
0%
100%
% of Puncta
26
13
0
0%
50%
100%
Position along Contact
D
E8
α-GFP α-PAR-3
HMR-1
HMP-2
HMP-1
AFD-1
PAR-6
PKC-3

**Figure S1**

**Supplemental Figure 1. Composition and localization of the LPC relative to intestinal anatomy.**

A) Cartoons of dorsal (top) and side (bottom) views of intestinal polarization from E8 to E16. Light grey E cells are destined for the ventral (‘V’) surface, yellow arrowheads signify sister cells. B) Dorsal view live imaging of endogenously GFP-tagged PKC-3 localization in a subset of cells in an ‘early’ vs ‘late’ E8 intestinal primordium. Images are single optical sections. At early E8, all cells occupy the same plane. At late E8, the contact between new dorsal neighbors (cells 1 and 3) lacks LPCs while in a ventral view (red), cell 2 and associated LPCs are visible (yellow arrowheads). C) LPC placement along the length of a homotypic contact (mean = 57%; n=64 puncta, 13 embryos). D) Dorsal view of E8 intestinal primordia (white dotted outline) in fixed embryos stained with indicated endogenously GFP-tagged protein (α-GFP (green)) and PAR-3 (α-PAR-3 (magenta)). Scale bars=5 μm (B, D) or 1 μm (C, D insets).

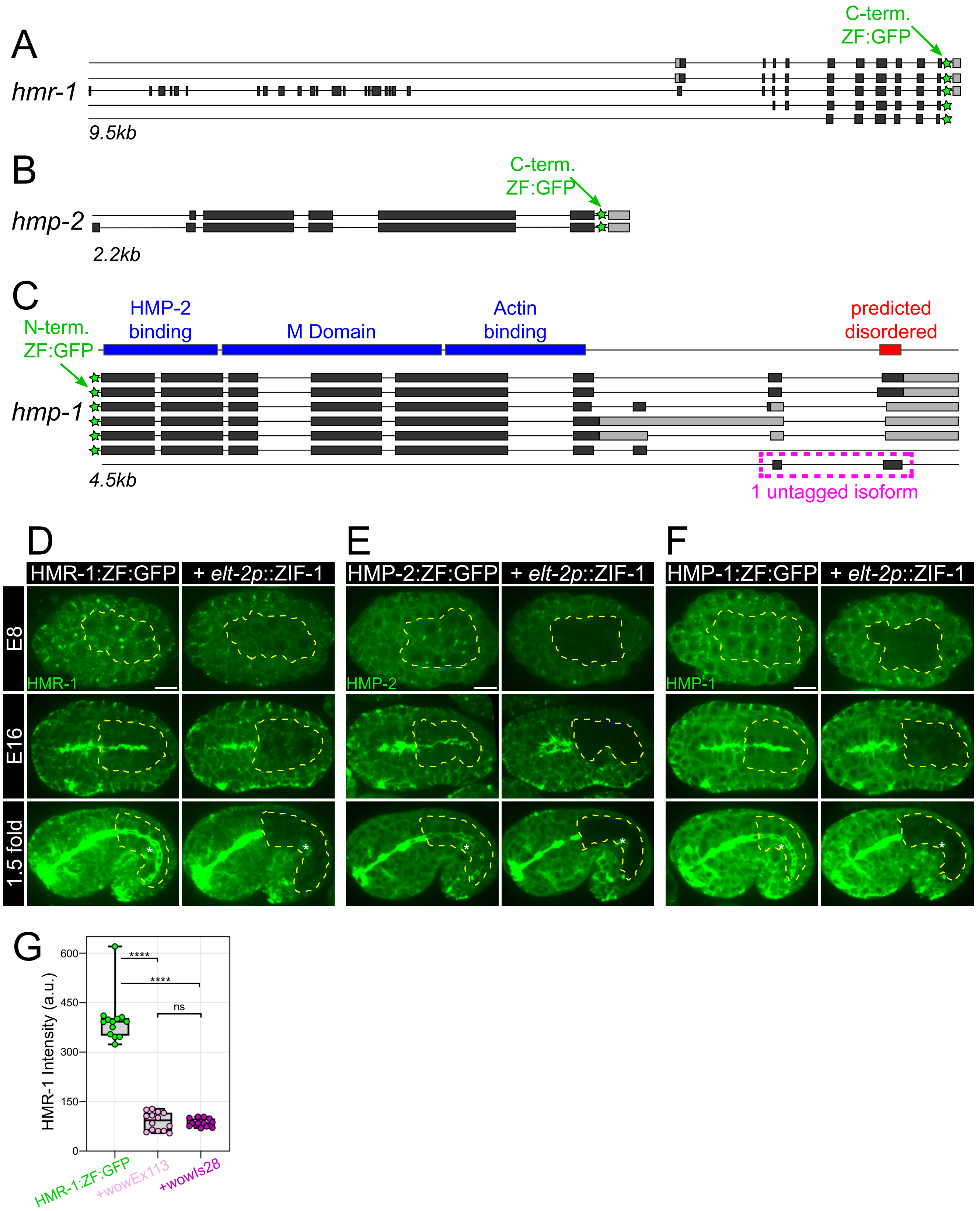

A
C-term.
ZF:GFP
hmr-1
9.5kb
B
C-term.
ZF:GFP
hmp-2
2.2kb
C
HMP-2
binding
M Domain
Actin
binding
predicted
disordered
N-term.
ZF:GFP
hmp-1
4.5kb
1 untagged isoform
D
HMR-1:ZF:GFP
+ elt-2p::ZIF-1
E
HMP-2:ZF:GFP
+ elt-2p::ZIF-1
F
HMP-1:ZF:GFP
+ elt-2p::ZIF-1
E8
E16
1.5 fold
HMR-1
HMP-2
HMP-1
G
HMR-1 Intensity (a.u.)
600
450
300
150
0
****
****
ns
HMR-1:ZF:GFP
+wowEx113
+wowIs28


**Figure S2**

**Supplemental Figure 2. Tissue-specific degradation of the cadherin/catenin complex.**

A-C) Known isoforms of *hmr-1*, *hmp-2*, and *hmp-1*, with *zf::gfp* tag location (green). C) Blue: Functional domains of HMP-1 protein, which are all absent from 1 untagged isoform (magenta) that only encodes a disordered region (red) and is predicted to be nonfunctional. D-F) Endogenously ZF::GFP-tagged proteins with (right) or without (left) intestinal ZIF-1 expression under the *elt-2* promoter (HMR-1:ZF:GFP, E8: n=16, E16: n=22, 1.5 fold: n=24; HMR-1:ZF:GFP+elt-2*p*:ZIF-1, E8: n=13, E16: n=21, 1.5 fold: n=21; HMP-2:ZF:GFP, E8: n=11, E16: n=13, 1.5 fold: n=13; HMP-2:ZF:GFP+elt-2*p*:ZIF-1, E8: n=6, E16: n=13, 1.5 fold: n=12; HMP-1:ZF:GFP, E8: n=18, E16: n=21, 1.5 fold: n=21; HMP-1:ZF:GFP+elt-2*p*:ZIF-1, E8: n=19, E16: n=29, 1.5 fold: n=27). Intestines outlined in dotted yellow lines. White asterisks = germ cells. G) HMR-1 intensity when undegraded (green) or degraded with two different *zif-1* alleles (see methods) ****$p<0.0001$. Scale bars=5 μm.

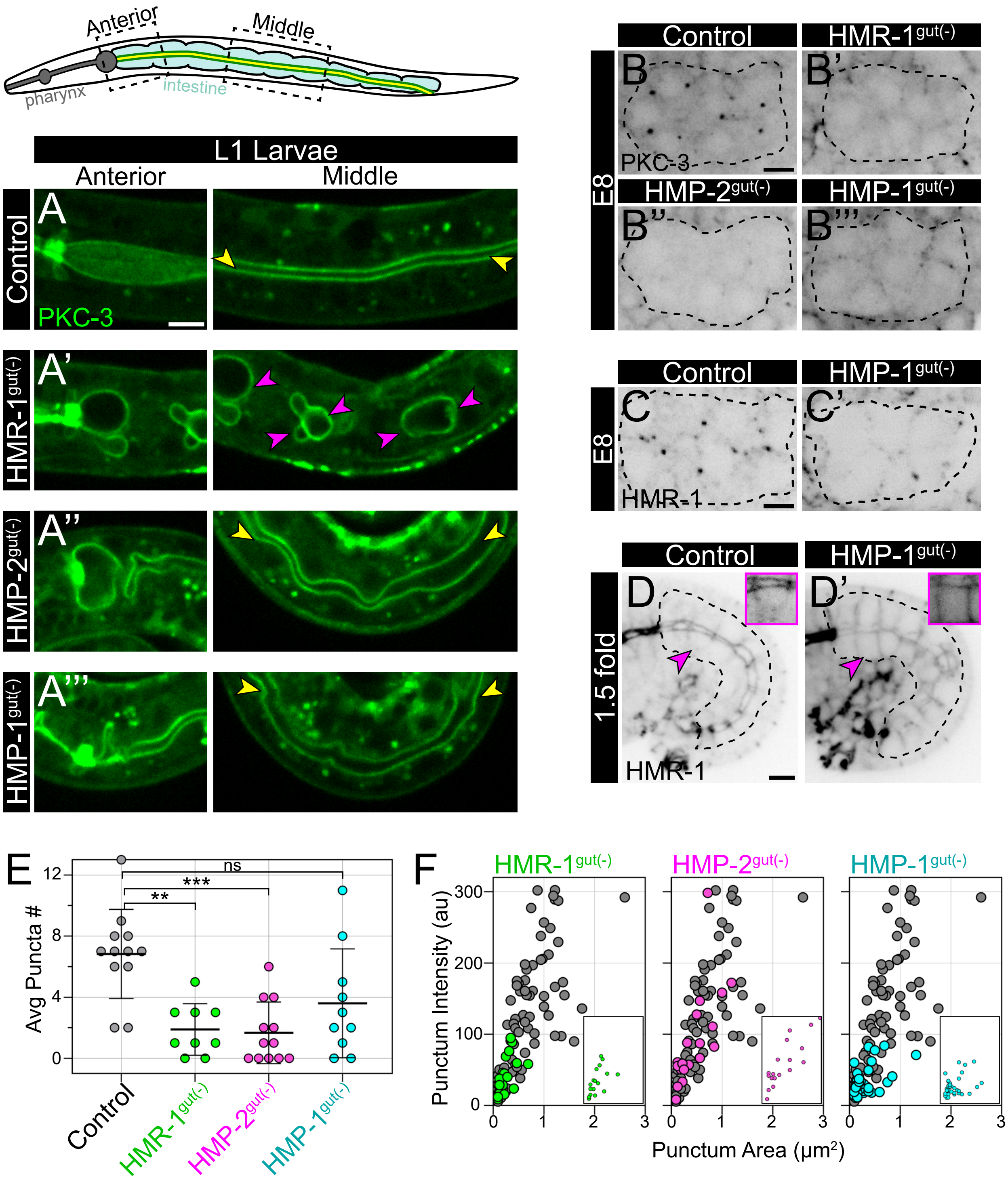

Anterior
Middle
pharynx
intestine
L1 Larvae
Anterior
Middle
Control
A
PKC-3
HMR-1gut(-)
A'
HMP-2gut(-)
A''
HMP-1gut(-)
A'''
Control
HMR-1gut(-)
E8
B
B'
PKC-3
HMP-2gut(-)
HMP-1gut(-)
B''
B'''
Control
HMP-1gut(-)
E8
C
C'
HMR-1
Control
HMP-1gut(-)
1.5 fold
D
D'
HMR-1
E
ns
***
**
Avg Puncta #
12
8
4
0
Control
HMR-1gut(-)
HMP-2gut(-)
HMP-1gut(-)
F
HMR-1gut(-)
HMP-2gut(-)
HMP-1gut(-)
Punctum Intensity (au)
300
200
100
0
0
1
2
3
Punctum Area (µm2)


**Figure S3**

**Supplemental Figure 3. Phenotypes differ between HMR-1 depletion and HMP-2 or HMP-1 depletion.**

A-A''') (Top) Cartoon of L1 larvae showing pharynx (dark grey) and intestinal cells (light blue), apical surfaces (green), and lumen (yellow). Boxes indicate regions shown in images below. (Bottom) Endogenously GFP tagged PKC-3 localization in two regions of the intestine (shown as single slices) in Control, HMR-1$^{gut(-)}$, HMP-2$^{gut(-)}$, and HMP-1$^{gut(-)}$ L1 larvae. A continuous lumen (yellow arrowhead) was observed in n = 17/17; 0/20; 14/14; 6/8, respectively. Note the cystic discontinuities (magenta arrowheads) in HMR-1$^{gut(-)}$ larvae. B-C') Dorsal view live imaging in E8 intestinal primordium expressing indicated endogenously GFP tagged protein. B-B''') Control (n=12), HMR-1$^{gut(-)}$ (n=9), HMP-2$^{gut(-)}$ (n=12), or HMP-1$^{gut(-)}$ (n=10). C-C') Control (n=13) or HMP-1$^{gut(-)}$ (n=9). D-D') Side view of E20-stage intestinal primordium in 1.5 fold stage embryos. Endogenous GFP-tagged HMR-1 in Control (n=4) or HMP-1$^{gut(-)}$ (n=8). Magenta insets show one E cell at high contrast. E) Average number of puncta per primordium in each genotype. One dot represents one embryo. ***$p<0.001$, **$p<0.01$. F) Puncta intensity vs. area in each indicated depletion background, with Control data behind in grey. Insets show tightly clustered points in bottom left of each graph plotted on a smaller scale. Scale bars=5 μm.

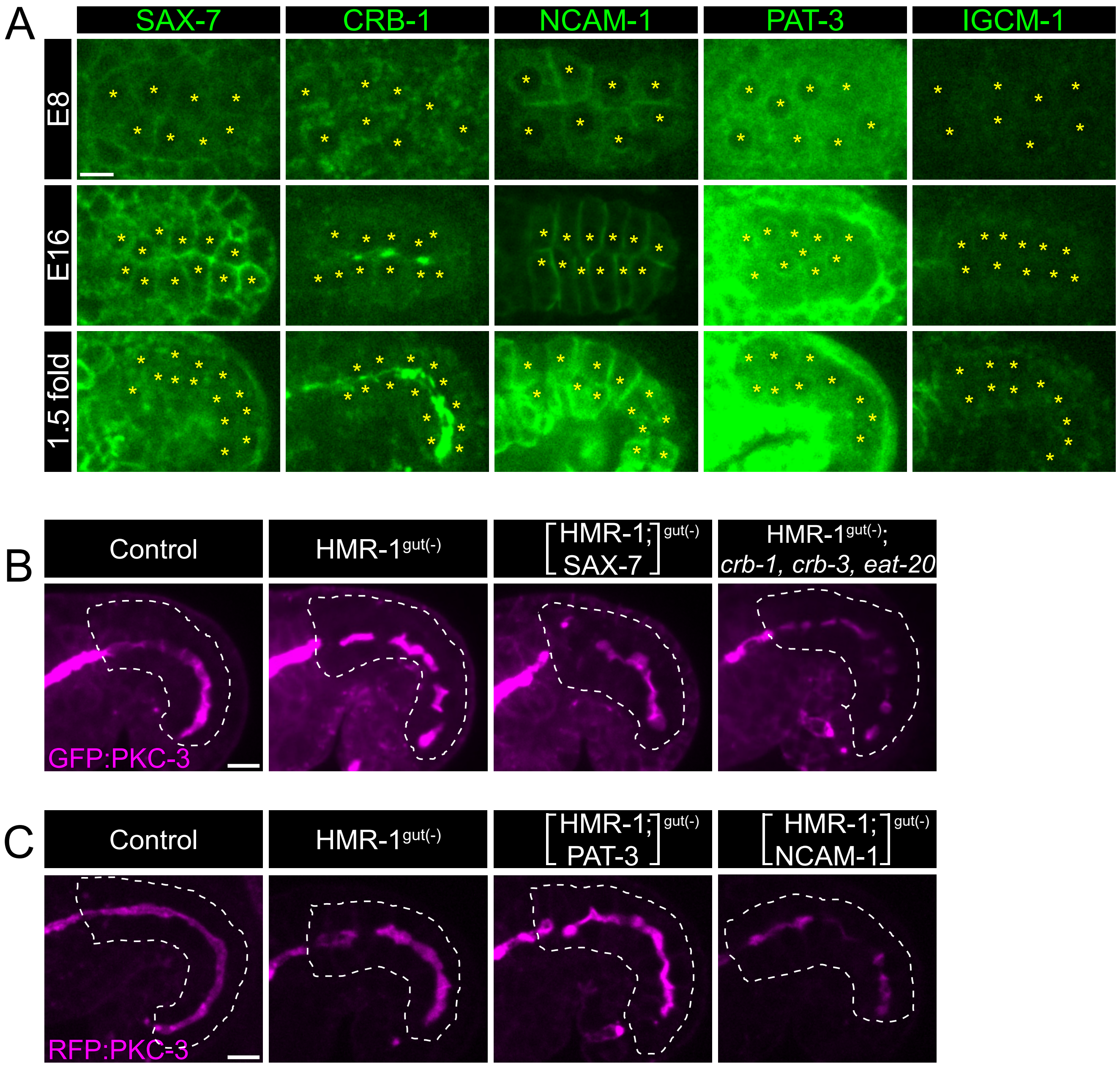

A
SAX-7
CRB-1
NCAM-1
PAT-3
IGCM-1
E8
E16
1.5 fold
B
Control
HMR-1gut(-)
[HMR-1; SAX-7]gut(-)
HMR-1gut(-); crb-1, crb-3, eat-20
GFP:PKC-3
C
Control
HMR-1gut(-)
[HMR-1; PAT-3]gut(-)
[HMR-1; NCAM-1]gut(-)
RFP:PKC-3


**Figure S4**

**Supplemental Figure 4. Candidate adhesion proteins fail to explain symmetry breaking.** A) Localization of indicated endogenously GFP-tagged proteins in E8, E16, and 1.5 fold stage intestinal primordia in live embryos, shown as single slices. E8 and E16 stages are dorsal views, while 1.5 fold stages are side views. Yellow asterisks highlight E cell nuclei. B-C) Live, side view images of Comma-stage embryos expressing endogenously tagged GFP::PKC-3 (B) or mScarlet::PKC-3 (C) in Control (B: n=10; C: n= 9), HMR-1$^{gut(-)}$ (B: n=10; C: n=5), or co-depletion of HMR-1 and SAX-7 ([HMR-1;SAX-7]$^{gut(-)}$, n=10), the Crumbs complex (HMR-1$^{gut(-)}$; *crb-1(0) crb-3(0) eat-20(0),* n=5), PAT-3 (([HMR-1;PAT-3]$^{gut(-)}$, n=7), or NCAM-1 (([HMR-1;NCAM-1]$^{gut(-)}$, n=5). All co-depletions achieved with ZF-tagging except where null mutations of the *crb-1, crb-3,* and *eat-20* were used. White dotted lines = intestine outline. Scale bars=5 µm.

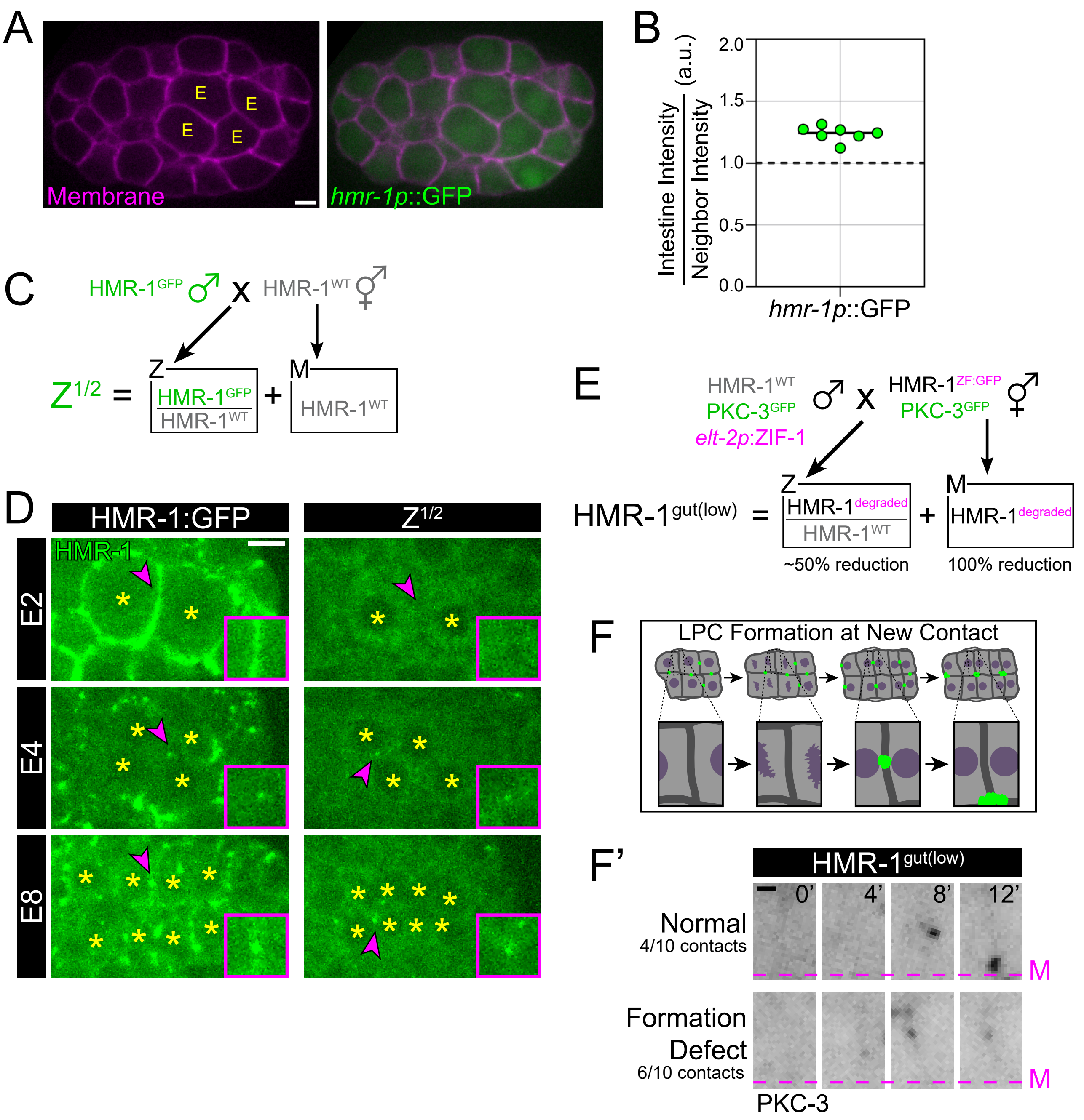
A
E
E
E
E
Membrane
hmr-1p::GFP
B
Intestine Intensity / Neighbor Intensity (a.u.)
2.0
1.5
1.0
0.5
0.0
hmr-1p::GFP
C
HMR-1GFP ♂ x HMR-1WT ⚥
Z1/2 = Z HMR-1GFP / HMR-1WT + M HMR-1WT
D
HMR-1:GFP
Z1/2
HMR-1
E2
E4
E8
E
HMR-1WT
PKC-3GFP
elt-2p:ZIF-1
♂ x
HMR-1ZF:GFP
PKC-3GFP
⚥
HMR-1gut(low) = Z HMR-1degraded / HMR-1WT + M HMR-1degraded
~50% reduction
100% reduction
F
LPC Formation at New Contact
F'
HMR-1gut(low)
0'
4'
8'
12'
Normal
4/10 contacts
Formation Defect
6/10 contacts
M
M
PKC-3

**Figure S5**

**Supplemental Figure 5. Intestinal HMR-1 levels play a supportive but nonessential role in LPC formation.**

A) Dorsal view of live E4 stage embryo expressing *hmr-1p*::GFP transcriptional reporter (green) and membrane::mCherry (magenta). Intestinal cells (‘E’) are indicated. B) Ratio of GFP signal inside E4 cells vs immediate neighbors is significantly different than 1 ($p<0.05$). C) Genetic scheme to achieve labeling of only zygotic HMR-1 (50% of all zygotic labeled, ‘$Z^{1/2}$’). D) Comparison of maternal and zygotic HMR-1::GFP (left) vs $Z^{1/2}$ labeling. Dorsal views of the intestinal primordium from indicated stages in live embryos. Yellow asterisks = E nuclei. Insets show region indicated by magenta arrowhead. Note the absence of zygotic HMR-1 at E2 but presence in foci at E4. E) Genetic scheme to achieve intestines with partial HMR-1 reduction (‘gut(low)’). F) Cartoon of LPC formation at the new cell/cell contact as two of the E cells move to the ventral tier at E8 (see Figure S1A) and the subsequent migration of these LPCs to the midline at E16. F’) Dorsal view live timelapse imaging of endogenously GFP tagged PKC-3 in the region depicted in (F). Defective LPC assembly or migration in 6/10 contacts from HMR-$1^{gut(less)}$ embryos. ‘M’ = midline, n=5 embryos. Scale bars=5 μm (A, D) or 1 μm (F’).

**Table S1.**

| Gene | Ortholog | Reason for testing | Onset of expression | Intestinal localization | Intestinal polarity phenotype |
|---|---|---|---|---|---|
| *sax-7* | L1CAM | Redundant with HMR-1 in early *C. elegans* embryo (Grana et al, 2010) | Before E | At all contacts | None |
| *crb-1, crb-3, eat-20* | Crumbs | Crumbs anisotropy polarizes myosin in *Drosophila* salivary gland (Röper, 2012) | E16 (CRB-1) | Apical | none |
| *ncam-1* | NCAM | RNAseq shows high expression in early *C. elegans* intestine (Hashimshony et al, 2012) | E8 | Intestinal membranes | none |
| *pat-3* | β-integrin | Required to polarize the *C. elegans* pharynx (Rasmussen et al, 2012) | E8 - E16 | Basal | none |
| *igcm-1* | Echinoid (*D. mel.)* | Echinoid polarizes myosin in *Drosophila* epidermis and cooperates with DE-cad (Wei et al, 2005; Laplante and Nilson, 2011) | E20 (~2-fold) | Basolateral | *not tested* |

### Table S2.

| Edit | Primer Sequences | | Plasmid |
|---|---|---|---|
| *wow152* | sgRNA | TTCAGATAAGACGTTCTTGgttttagagctagaaatagcaagt | pVN44 |
| *[crb-1::gfp]* | 5' HA F | ttgtaaaacgacggccagtcgccggcaTCACACAGAATCGAATGGCGGTT | pVN45 |
| | 5' HA R | CATCGATGCTCCTGAGGCTCCCGATGCTCCGATAAGTCGCTCCTGAGGTGGAA | |
| | 3' HA F | CGTGATTACAAGGATGACGATGACAAGAGATGAaccctgaatacctgaattatttg | |
| | 3' HA R | ggaaacagctatgaccatgttatcgatttcTGAGCGAGTTCAACATTTCACTT | |
| *wow138* | sgRNA | TTTTCAGAATGCCTGCGAAgttttagagctagaaatagcaagt | pVN35 |
| *[zf::gfp::hmp-1]* | 5' HA F | ttgtaaaacgacggccagtcgccggcaCTCACTCGCAAATACCTCAA | pVN34 |
| | 5' HA R | ACAAAGTCGCGTTTTGTATTCTGTCGGCATactgaaaattaataaaattgaaaattc | |
| | 3' HA F | CGTGATTACAAGGATGACGATGACAAGAGACCGGCCAATGGCAATTCTC | |
| | 3' HA R | tcacacaggaaacagctatgaccatgttatGGCTCGTCGTCGAACAGTGA | |
| *wow99* | sgRNA | ATGAAAAAGAAGGAACACGgttttagagctagaaatagcaagt | pVN23 |
| *[sax-7::zf::gfp]* | 5' HA F | ttgtaaaacgacggccagtcgccggcaTGTTCGTGTTGCTGCTCTTC | pVN24 |
| | 5' HA R | CATCGATGCTCCTGAGGCTCCCGATGCTCCGACAAACGTCGACGTTGATCCT | |
| | 3' HA F | CGTGATTACAAGGATGACGATGACAAGAGATAGaagactatttacgtgttccttcttt | |
| | 3' HA R | ggaaacagctatgaccatgttatcgatttctccgcatattactgggatga | |
| *wow108* | sgRNA | AGATAAATTAATTACAAATgttttagagctagaaatagcaagt | pVN31 |
| *[hmp-2::zf::gfp]* | 5' HA F | ttgtaaaacgacggccagtcgccggcaAGCCACCACGGATTTAAGATATTG | pVN30 |
| | 5' HA R | CATCGATGCTCCTGAGGCTCCCGATGCTCCCAAATCAGTATCGTACCAATTGTGA | |
| | 3' HA F | CGTGATTACAAGGATGACGATGACAAGAGATAAttaatttatcttctcgtatttttgcat | |
| | 3' HA R | ggaaacagctatgaccatgttatcgatttcTTTTTCCAATAAATCCTCCGTACAA | |
| *wow110* | sgRNA | TTAAAAATCCAGTATACGCgttttagagctagaaatagcaagt | pVN33 |
| *[pat-3::zf::gfp]* | 5' HA F | ttgtaaaacgacggccagtcgccggcaCAACGAAACTACACCCTGCCA | pVN32 |
| | 5' HA R | CATCGATGCTCCTGAGGCTCCCGATGCTCCGTTGGCTTTTCCAGCATAGACAGGAT TTTT | |
| | 3' HA F | CGTGATTACAAGGATGACGATGACAAGAGATAAatagtttttatccttatattttaataattttcc | |
| | 3' HA R | ggaaacagctatgaccatgttatcgatttcCTTTCTGTGCACTTGGATGC | |
| *wow107* | sgRNA | GAGAGATTTGGAGTCCGGAGTTTTAGAGCTAGAAATAGCAAGT | pVN29 |
| *[ncam-1::zf::gfp]* | 5' HA F | TTGTAAAACGACGGCCAGTCGCCGGCAACACTTTTGAAGCGAGAAGACT | pVN28 |
| *(internal)* | 5' HA R | CATCGATGCTCCTGAGGCTCCCGATGCTCCTTTAGCACCAGCGTTCTTTC | |
| | 3' HA F | CGTGATTACAAGGATGACGATGACAAGAGACAGCGAGATCTTGAGTCGGGAAG | |
| | 3' HA R | GGAAACAGCTATGACCATGTTATCGATTTCTGCAATATGAAACATGGAAACCAA | |
| *wow167* | sgRNA | AATGAAAGAAGTTTGTGAGgttttagagctagaaatagcaagt | pVN63 |
| *[igcm-1::zf::gfp]* | 5' HA F | ttgtaaaacgacggccagtcgccggcaTGATGCAAAGTCGACGAAAC | pVN64 |
| | 5' HA R | CATCGATGCTCCTGAGGCTCCCGATGCTCCAACAATAATCTCTCGAACGATTC | |
| | 3' HA F | CGTGATTACAAGGATGACGATGACAAGAGATAAttttgtcatatttttgtcattcttccaacc | |
| | 3' HA R | ggaaacagctatgaccatgttatcgatttcggtcggcataccatgatcg | |
